# Novel adaptation supports enhanced macrophage efferocytosis in limited-oxygen environments

**DOI:** 10.1101/2022.08.04.502838

**Authors:** Ya-Ting Wang, Alissa Trzeciak, Waleska Saitz Rojas, Pedro Saavedra, Yan-Ting Chen, Rachel Chirayil, Jon Iker Etchegaray, Christopher D. Lucas, Daniel J. Puleston, Kayvan R. Keshari, Justin S. A. Perry

## Abstract

Apoptotic cell clearance (efferocytosis), a process essential for organismal homeostasis, is performed by phagocytes that inhabit a wide range of environments, including physiologic hypoxia. Here, we find macrophages, the predominant tissue-resident phagocyte, display enhanced efferocytosis under prolonged (chronic) physiological hypoxia, characterized by increased internalization and accelerated degradation of apoptotic cells. Analysis of mRNA and protein programs revealed that chronic physiological hypoxia induces two distinct but complimentary states in macrophages. The first, ‘primed’ state consists of concomitant induction of transcriptional and translational programs broadly associated with metabolism in apoptotic cell-naïve macrophages that persist during efferocytosis. The second, ‘poised’ state consists of transcription, but not translation, of phagocyte function programs in apoptotic cell-naïve macrophages that are subsequently translated during efferocytosis. Importantly, we discovered that both states are necessary for enhanced continual efferocytosis. Mechanistically, we find that one such ‘primed’ state consists of the efficient flux of glucose into a noncanonical pentose phosphate pathway (PPP) loop, whereby PPP-derived intermediates cycle back through the PPP to enhance production of NADPH. Furthermore, we found that PPP-derived NADPH directly supports enhanced continual efferocytosis under chronic physiological hypoxia via its role in phagolysosomal maturation and maintenance of cellular redox homeostasis. Thus, macrophages residing under chronic physiological hypoxia adopt states that both support cell fitness and ensure ability to perform essential homeostatic functions rapidly and safely.

**Highlights:**

- Macrophages residing in chronic physiological hypoxia have enhanced apoptotic cell uptake and degradation
- Chronic physiological hypoxia induces both primed and poised states in macrophages
- Both primed and poised state programs directly support enhanced continual efferocytosis
- A noncanonical PPP loop, a unique primed state, directly supports enhanced efferocytosis and maintains redox homeostasis

## Introduction

Efferocytosis, the phagocytic clearance of apoptotic cells, is indispensable for organismal homeostasis (Boada-Romero et al., 2020; Doran et al., 2020; Rothlin et al., 2020). Efferocytosis occurs in all major tissues and organs, ensuring that disparate processes such as barrier epithelial cell recycling, removal of spent neutrophils and red blood cells, elimination of apoptotic neurons, and clearance of negatively-selected thymocytes are executed rapidly and safely (Elliott and Ravichandran, 2016; Henson, 2017; Morioka et al., 2019). These clearance processes are performed by various types of phagocytes, especially by tissue-resident macrophages (TRMs; (Penberthy et al., 2018; Zago et al., 2021)). At homeostasis, each tissue exhibits a substantial rate of cellular turnover, collectively amounting to ∼1-2% body mass each day (Sender and Milo, 2021). As the body’s main professional phagocyte, TRMs shoulder the brunt of this massive burden (Trzeciak et al., 2021).

TRMs are generally long-lived cells that seed a tissue early during development and often reside in a tissue throughout the organism’s lifespan (Amit et al., 2016; Guilliams and Svedberg, 2021; Guilliams et al., 2020; Okabe and Medzhitov, 2016). It is now clear that TRMs adapt to their unique and often harsh tissue environment to perform their core functions (Amit et al., 2016; Blériot et al., 2020; Guilliams et al., 2020), including residing in tissues with a relative dearth of oxygen availability and exposure to tissue-specific debris and metabolites. For instance, the tissues that feature the most cell turnover (bone marrow, spleen, thymus; (Breed et al., 2019; Sender and Milo, 2021; Stritesky et al., 2013)) are also tissues with the lowest oxygen availability (∼1% O_2_, termed physiological hypoxia; (Carreau et al., 2011; Norris et al., 2019; Trayhurn, 2019)). TRMs, then, must both adapt to the tissue environment in which they reside and cope with the continuous influx of internalized biological material. Recent work has illustrated mechanisms by which phagocytes sense and respond to internalized apoptotic cells (Morioka et al., 2018; Perry et al., 2019; Wang et al., 2017; Yurdagul et al., 2020; Zhang et al., 2019), however, how tissue environment factors, such as oxygen availability, inform the ability to perform efferocytosis remains unknown.

Here, we made the striking discovery that exposure to prolonged (‘chronic’) physiological hypoxia, similar to that experienced by several TRM populations, resulted in increased internalization and accelerated degradation of apoptotic cells. We found that chronic exposure to physiological hypoxia induced two distinct but complimentary states. One state, which we term ‘primed’, generally consists of simultaneous transcriptional and translational induction or suppression of metabolic programs in apoptotic cell-naïve macrophages that remain induced or suppressed during efferocytosis. The other state, which we term ‘poised’, generally consists of transcription, without concomitant translation, of phagocytic functional programs in apoptotic cell-naïve macrophages that are instead translated during efferocytosis. Importantly, we discovered that both states are necessary for enhanced continual efferocytosis. Subsequent exploration of primed state metabolic programs revealed that macrophages exposed to chronic physiological hypoxia switch to the efficient utilization of glucose to generate NADPH via a noncanonical pentose phosphate pathway (PPP) loop that features recycling of PPP-derived intermediates back through the oxidative PPP. The PPP-dependent generation of NADPH prior to efferocytosis served to both support enhanced internalization and degradation of apoptotic cells via phagolysosomal maturation and protect phagocytes from runaway oxidative stress. These studies reveal that local tissue environment, in this case physiological hypoxia, programs distinct states in macrophages that simultaneously support cell fitness and ensure the ability to perform critical homeostatic functions, such as efferocytosis.

## Results

### Efferocytosis is enhanced under prolonged (‘chronic’) physiological hypoxia

Professional phagocytes, such as tissue-resident macrophages, often reside for long periods of time in environments with extremely low (e.g., 1%) oxygen availability (physiological hypoxia; (Blériot et al., 2020; Guilliams et al., 2020)). Despite this relative dearth of oxygen, phagocytes must clear millions of apoptotic cells, a homeostatic process known as ‘efferocytosis’. To address the effect of prolonged (‘chronic’) physiological hypoxia on efferocytosis, we optimized a system and protocols to continually manipulate and culture primary professional phagocytes (macrophages) in low (1%) oxygen. Strikingly, chronic physiological hypoxia-conditioned macrophages engulfed significantly more apoptotic cells than both macrophages cultured under atmospheric (‘standard’) oxygen levels (21%) and macrophages exposed to ‘acute’ (3h) hypoxia (Figure 1A). Chronic physiological hypoxia-conditioned macrophages engulfed significantly more apoptotic cells irrespective of the size of the target cell, the type of target cell, the method of cell death induction, or the transformation status of the target cell (Figure 1B; Figure S1A-D). Furthermore, macrophages engulfed significantly more apoptotic cells regardless of whether they were initially differentiated in standard oxygen conditions or differentiated under physiological hypoxia prior to conditioning (Figure 1C).

**Figure 1:**
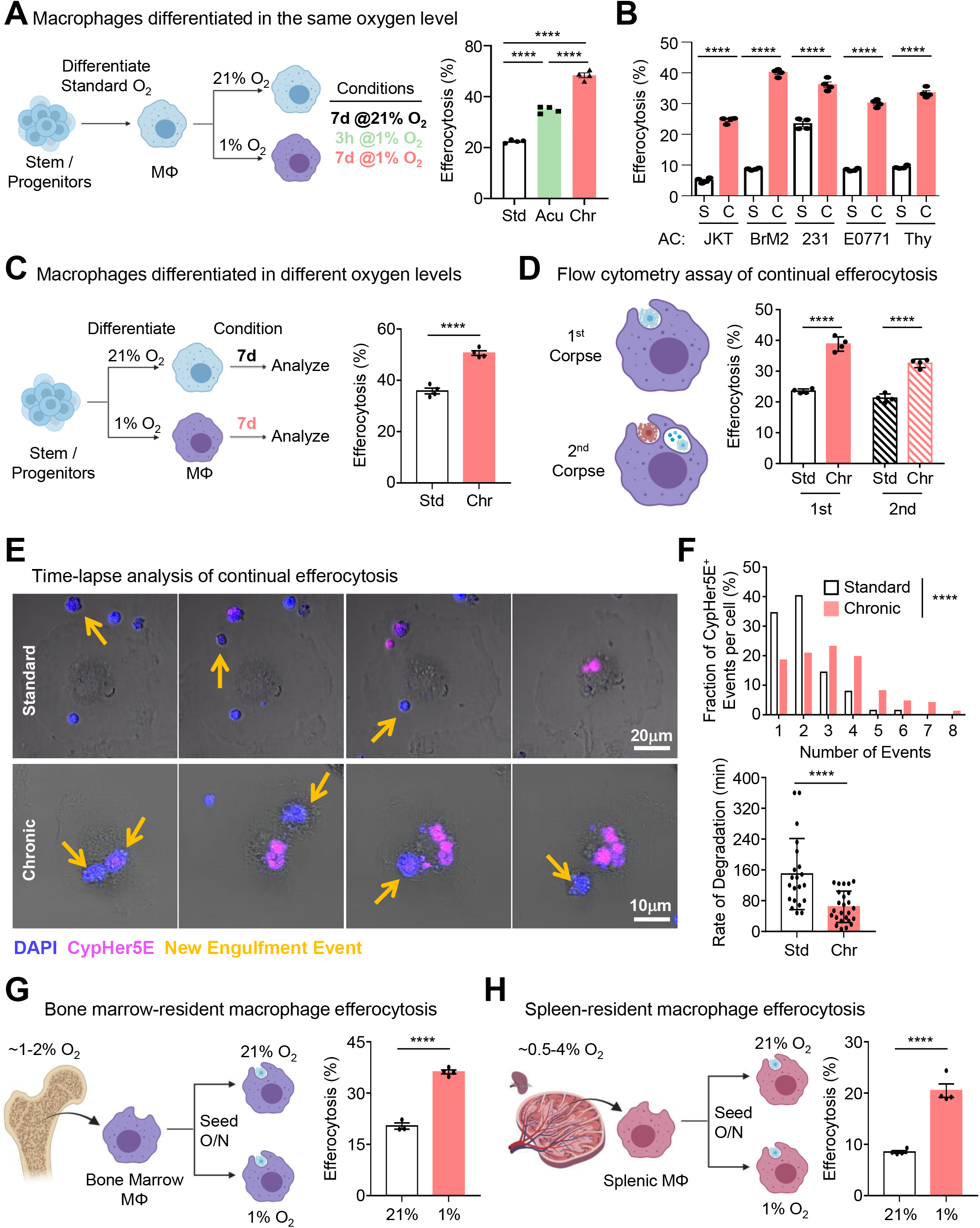
Efferocytosis is enhanced under prolonged (‘chronic’) physiological hypoxia. (A) <u>Efferocytosis by macrophages differentiated in the same oxygen environment, conditioned in</u> <u>standard oxygen or physiological hypoxia.</u> (Left) ER-Hoxb8 macrophage progenitors were differentiated under standard (21%) O_2_ for 6 days. The macrophages were then replated and conditioned for 7d in 21% O_2_ (Std), 1% O_2_ (Chr) or 7 days in 21% O_2_ followed by 3h in 1% O_2_ (Acu). (Right) Conditioned macrophages were co-cultured with CypHer5E-labeled apoptotic MDA-MB-231 cells at a 1:1 ratio for 1h. Efferocytosis was assessed via FACS. Data are from four independent experiments. Data are shown as mean ± SEM. ****p < .0001. (B) <u>Efferocytosis by standard oxygen- or physiological hypoxia-conditioned macrophages of</u> <u>different apoptotic targets.</u> Standard or chronic physiological hypoxia-conditioned macrophages as in (A) were co-cultured with CypHer5E-labeled apoptotic cells at a 1:1 ratio. The following target types were used: Jurkat T cells, BrM2s, MDA-MD-231s, E0771s, and primary thymocytes. Efferocytosis was assessed via FACS. Data are from three independent experiments. Data are shown as mean ± SEM. ****p < .0001. (C) <u>Efferocytosis by macrophages differentiated in different oxygen levels.</u> (Left) ER-Hoxb8 progenitors were differentiated under standard O_2_ (21%) or physiological hypoxia (1%) for 6d. Then, macrophages were replated and maintained in either 21% O_2_ (Std) or 1% O_2_ (Chr) for 7d. (Right) Macrophages were co-cultured with CypHer5E-labeled apoptotic MDA-MB-231 at a 1:1 ratio for 1h. Efferocytosis was assessed via FACS. Data are from three independent experiments. Data are shown as mean ± SEM. ****p < .0001. (D) <u>Chronic hypoxic conditioning promotes continual efferocytosis by macrophages.</u> Conditioned macrophages as in (A) were first co-cultured with CFSE-labeled apoptotic MDA-MB-231 cells at a 1:1 ratio for 1h. Unengulfed apoptotic cells were removed by BMDM media wash (3x). Cells were rested for 1.5h, then co-cultured with CypHer5E-labeled apoptotic MDA-MB-231 cells at a 1:1 ratio for 1h and analyzed via FACS. Data are from three independent experiments. Data are shown as mean ± SEM. ****p < .0001. (E, F) <u>Time-lapse microscopy analysis of continual efferocytosis.</u> (E) Macrophages were differentiated as in (A), replated in glass-bottom dishes, then conditioned in standard O_2_ (21%) or physiological hypoxia (1%) for 7d. Macrophages were co-cultured with DAPI (blue)/CypHer5E (pink)-labeled apoptotic Jurkat T cells at a 1:1 ratio. Imaging was started ∼5-10min after the addition of apoptotic cells. Yellow arrows indicate newly engulfed apoptotic cells. (F) Quantification of the number of CypHer5E+ events (top) and the rate of degradation of engulfed apoptotic cells (bottom). For quantification, 115 efferocytotic macrophages from 8 standard oxygen scenes and 155 efferocytotic macrophages from 6 chronic physiological hypoxia scenes were analyzed. Data were binned as number of events per cell and presented as a fraction of 100%. For analysis of degradation rate, 21 (standard oxygen) and 25 (chronic physiological hypoxia) efferocytotic macrophages were analyzed. Time to degradation was defined as the time it takes to shrink an internalized corpse 50% after initial acidification (CypHer5E+). (G, H) <u>Low oxygen environment supports enhanced efferocytosis by splenic- and bone marrow-resident macrophages.</u> Bone marrow (G) or spleen (H) were harvested from 7-week-old female C57BL/6J mice. Cells were isolated in a hypoxia chamber at 1% oxygen and seeded in either 1% or 21% oxygen overnight. Adherent macrophages were then co-cultured with CypHer5E-labeled apoptotic MDA-MD-231 cells at a 1:1 ratio for 1h. Efferocytosis was assessed via FACS. Data are from four independent experiments. Data are shown as mean ± SEM. ****p < .0001.

Professional phagocytes are often responsible for engulfment of apoptotic cells in quick succession, a phenomenon known as ‘continual efferocytosis’, which protects against the manifestation of inflammatory disease, such as atherosclerosis (Park et al., 2011; Viaud et al., 2018; Wang et al., 2017; Yurdagul et al., 2020). Interestingly, chronic physiological hypoxia-conditioned macrophages also exhibited significantly higher levels of continual efferocytosis (Figure 1D). Despite the increased proportion of macrophages engulfing more apoptotic cells on a population level, our flow cytometric analysis of the geometric mean fluorescence intensity of apoptotic cell uptake (a surrogate of uptake events per cell) was similar between conditions (Figure S1E), suggesting that either 1) individual macrophages do not take up more corpses, or 2) chronic physiological hypoxia-conditioned macrophages engulf more apoptotic cells and digest engulfed apoptotic cells quicker on a per cell basis. To test these hypotheses, we performed time-lapse confocal microscopy of macrophages conditioned in standard oxygen or under chronic physiological hypoxia. Indeed, chronic physiological hypoxia-conditioned macrophages not only engulfed significantly more apoptotic cells on a per cell basis (∼60% engulfed 3 or more corpses per cell compared to 25% of macrophages cultured in standard oxygen) but also degraded engulfed apoptotic cells significantly faster (Figure 1E and 1F).

Multiple tissue-resident macrophage populations reside under physiological hypoxia. For instance, bone marrow-resident macrophages and macrophage subsets in the spleen chronically experience oxygen levels as low as ∼1% (Norris et al., 2019). To test the *in vivo* relevance of physiological hypoxia on efferocytosis, we isolated tissue-resident macrophages (TRMs) from either the bone marrow or spleen and seeded them in either standard or 1% oxygen (Figure 1G and 1H). Consistent with our *in vitro* conditioning studies, we found that both bone marrow and splenic TRMs exhibited higher efferocytosis when maintained in 1% oxygen compared to standard oxygen (Figure 1G and 1H; Figure S1F). Collectively, our data suggest that professional phagocytes better engulf and digest potentially dangerous apoptotic cells under prolonged physiological hypoxia.

### Characterization of macrophages under chronic physiological hypoxia

To investigate how macrophages adapt to chronic physiological hypoxia, we performed RNA sequencing (RNAseq) of primary macrophages cultured in standard oxygen (21%), exposed to acute hypoxia (1% for 3h), and exposed to prolonged physiological hypoxia (1% for 7d; Figure 2A). Analysis of chronic physiological hypoxia-conditioned macrophages revealed several differentially expressed transcriptional programs (Figure 2B). For instance, we observed significant downregulation of lipid and mitochondrial metabolism programs and upregulation of carbohydrate metabolism and hypoxia-responsive programs (Figure 2B), broadly consistent with a previous study of cancerous cells cultured under chronic hypoxia (Jain et al., 2020). Importantly, we observed several programs not previously associated with physiological hypoxia generally, nor chronic physiological hypoxia specifically. For instance, we observed downregulation of metabolic programs (amino acid metabolism, oxidative stress/redox biology, insulin signaling), cell biological processes (endocytosis, mitochondrial biology, autophagy, proapoptotic signaling, exocytosis, vesicular transport), immune cell function (pro-inflammatory processes, antigen presentation), ion transport, and calcium homeostasis (Figure 2B; Figure S2A and S2B). On the other hand, we observed upregulation of key homeostatic macrophage function programs (phagocytosis, anti-inflammatory processes, wound healing, ECM Biology, angiogenesis), cell biological processes (lysosome biology, ER Stress/UPR, actin/cytoskeleton, cell adhesion, cell migration, anti-apoptotic signaling), and putative pro-tumorigenic programs (Figure 2B; Figure S2A and S2B). Although the majority of the chronic physiological hypoxia-specific program was induced only after prolonged hypoxia, a fraction of the program was modestly, albeit non-significantly, induced in response to acute hypoxia (Chronic vs. Standard Only; Figure 2C). Because these genes continue to significantly increase after prolonged physiological hypoxia, we consider them as part of the chronic physiological hypoxia-induced program.

**Figure 2:**
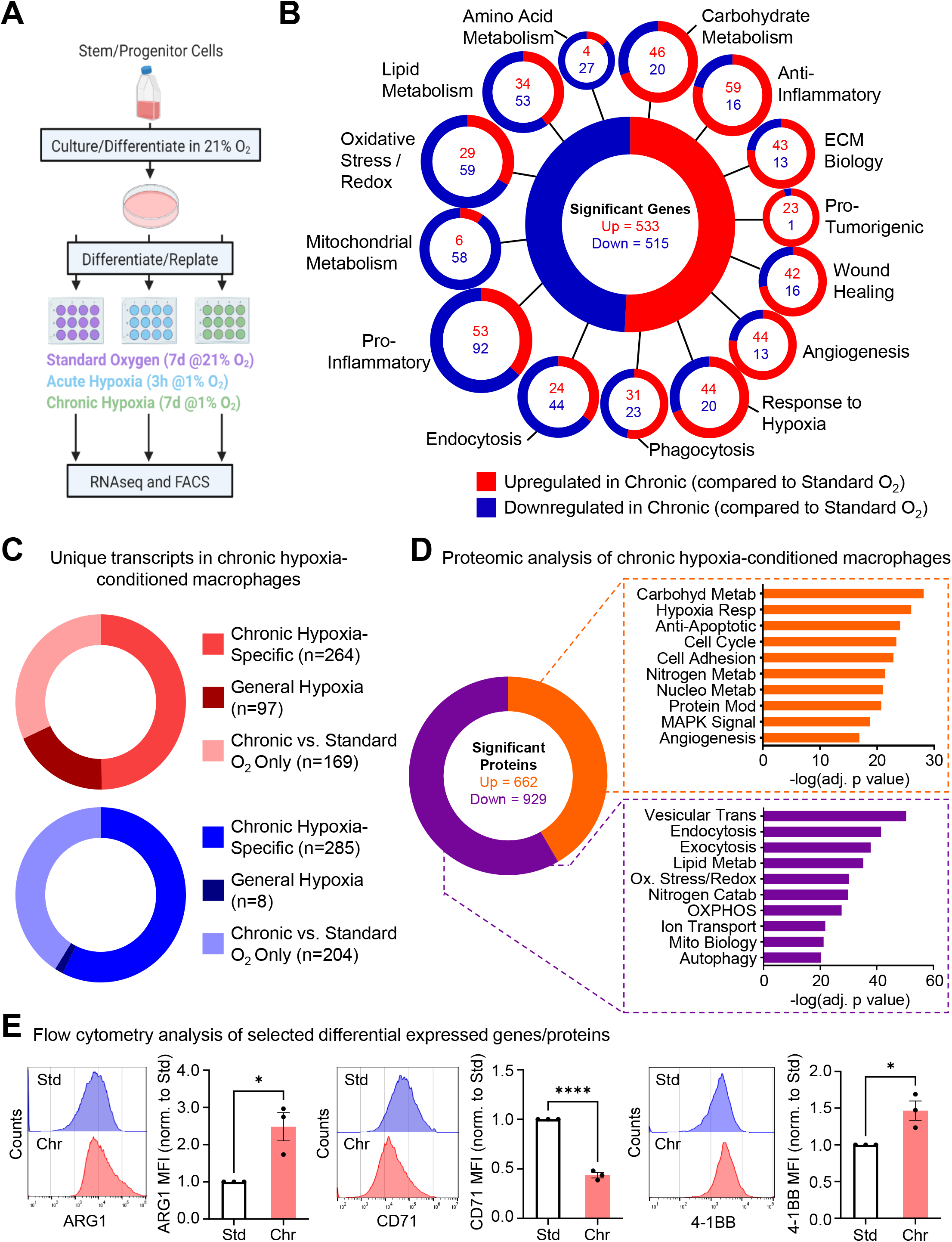
Characterization of macrophages under chronic physiological hypoxia. (A, B) <u>RNA sequencing analysis of macrophages conditioned in different oxygen environments</u> <u>reveals regulation of novel functional programs in chronic physiological hypoxia-conditioned</u> <u>macrophages.</u> (A) Schematic of method used for RNA sequencing analysis. ER-Hoxb8 progenitors were differentiated under standard (21%) oxygen for 6d. Macrophages were then replated and conditioned for 7d in standard (21%) oxygen (Std), for 7d in physiological hypoxia (1%; Chr), or in standard oxygen for 7d followed by 3h of physiological hypoxia (Acu). Macrophages were lysed in the conditioned oxygen environments for cell and mRNA extraction. (B) We detected significant changes in 1,048 genes in chronic hypoxia-conditioned macrophages relative to standard oxygen-conditioned macrophages. Differentially expressed genes were classified according to known or putative (based on family or sequence similarity) function. Genes were considered significant if they met a false discovery rate-adjusted p value < .05. Three independent experiments were analyzed for each condition. ECM, extracellular matrix. (C) <u>Identification of transcripts uniquely induced by chronic physiological hypoxia conditioning.</u> Analysis of data from (A) to specifically identify transcripts differentially regulated in chronic physiological hypoxia-conditioned macrophages compared to standard oxygen and acute physiological hypoxia (‘Chronic Hypoxia-Specific’), significantly differentially regulated under acute physiological hypoxia and maintained during chronic conditioning (‘General Hypoxia’), or transcripts that are only significantly differentially regulated in chronic physiological hypoxia-conditioned macrophages compared to standard oxygen (‘Chronic vs. Standard O_2_ Only’). (D) <u>Chronic physiological hypoxia-conditioned macrophages express unique proteins and induce</u> <u>or suppress distinct functional programs.</u> Experiments were performed as in (A) but instead analyzed via TMT-labeled mass spectrometry (proteomics). We detected significant changes in 1,591 proteins in chronic physiological hypoxia-conditioned macrophages relative to standard oxygen-conditioned macrophages. Differentially expressed proteins were classified according to known or putative (based on family or sequence similarity) function. Proteins were considered significant if they met a false discovery rate-adjusted p value < .05. Six independent experiments were analyzed for each condition. MAPK, mitogen-activated protein kinase. OXPHOS, oxidative phosphorylation. Pathway significance was determined using Fisher’s Exact Tests and presented as the negative log of the adjusted p value. (E) <u>Flow cytometric validation of selected differentially expressed genes/proteins.</u> Macrophages were conditioned as in (A), then harvested and analyzed for Arginase 1 (Arg1), CD71, and 4-1BB expression by flow cytometry. Shown are representative FACS histograms and normalized mean fluorescence intensity (MFI). Data represent three independent experiments and are shown as mean ± SEM. *p < 0.05. ****p < .0001.

In parallel, we performed proteomic analysis of chronic physiological hypoxia-conditioned macrophages. Our analysis revealed several differentially regulated programs that overlap with programs identified via RNAseq, especially programs involved in cellular metabolism (e.g., upregulation of carbohydrate metabolism, downregulation of lipid and mitochondrial metabolism; Figure 2D). We also observed upregulation of programs involved in nucleotide and nitrogen metabolism, and downregulation of a program involved in nitrogen catabolism. We subsequently validated representative targets fitting into one of three criteria: 1) a target that was upregulated in both RNAseq and proteomics analysis, Arginase 1 (ARG1, gene symbol *Arg1*), 2) a target that was downregulated in both RNAseq and proteomics analysis, transferrin receptor protein 1 (CD71, gene symbol *Tfrc*), and 3) a target that was upregulated in RNAseq but not detected via proteomics 4-1BB (CD137, gene symbol *Tnfrs9*) (Figure 2E). Taken together, these data indicate that macrophages induce unique gene and protein programs in response to chronic physiological hypoxia and that these adaptations support essential macrophage function in limited oxygen environments.

### Chronic physiological hypoxia induces states both primed and poised for efferocytosis

Beyond the differentially-regulated genes that we failed to detect signal for in our proteomics analysis, we also made the striking observation that a plurality of genes significantly differentially regulated by chronic hypoxia are detected by RNA sequencing but remain unchanged at the protein level (Figure S3A). One hypothesis is that macrophages in specific environments, such as a tissue with low oxygen, accumulate function-specific mRNAs that allow for rapid protein synthesis in response to performing a canonical function. To test this, we used a strategy for performing mass spectrometry analysis of proteins in efferocytotic macrophages while excluding contaminating peptides from engulfed apoptotic cells. Specifically, we labeled live target cells with ^13^C-lysine for >7 generations to stable isotope-label amino acids prior to induction of apoptosis and co-culture with macrophages (diagram in Figure 3A; global analyses in Figure S3B). Analysis of the core chronic physiological hypoxia-induced mRNA program revealed two distinct patterns of protein expression dynamics. <u>First</u>, we observed transcriptional programs that were also significantly differentially expressed at the protein level in apoptotic cell-naïve macrophages, which typically remained significantly increased/decreased (or increased/decreased further) in efferocytotic macrophages (‘Primed’ Programs in Figure 3A; Figure S4, PFKL/*Pfkl* as a representative example in Figure 3B). This cluster consisted primarily of cellular metabolism programs, such as carbohydrate metabolism, lipid metabolism, and mitochondrial metabolism. <u>Second</u>, and far more surprisingly, we observed programs differentially regulated at the mRNA level but not at the protein level in apoptotic cell-naïve macrophages, that were subsequently significantly differentially regulated at the protein level in efferocytotic macrophages (‘Poised’ Programs in Figure 3A; Figure S4, SAMD9L/*Samd9l* as a representative example in Figure 3B). This cluster consisted primarily of macrophage function programs, including phagocytosis, ECM biology, and cell adhesion. Thus, chronic physiological hypoxia induces two distinct but complementary states in macrophages. The first ‘primed’ state responds immediately by coupling transcription and translation to adapt to the local environment and prime apoptotic cell-naïve macrophages for efferocytosis. The second ‘poised’ state features transcriptional changes without commensurate protein translation in apoptotic cell-naïve macrophages, instead poising apoptotic cell-naïve macrophages with function-specific transcripts that are subsequently translated *during* efferocytosis.

**Figure 3:**
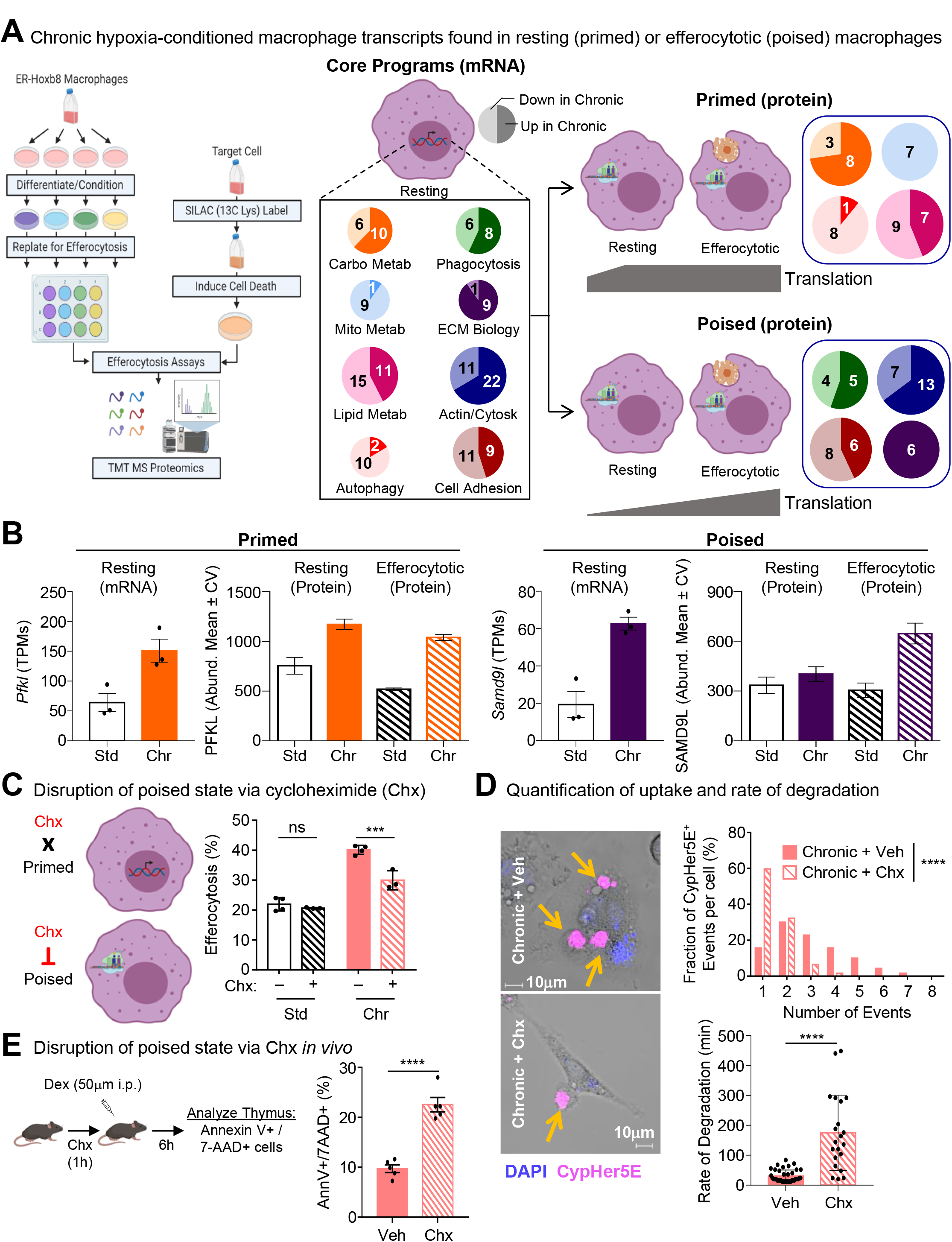
Chronic hypoxia induces states both primed and poised for efferocytosis. (A) <u>Global proteomic analysis of naive and efferocytotic macrophages conditioned in standard</u> <u>oxygen or chronic physiological hypoxia reveals ‘primed’ and ‘poised’ states.</u> (Left) ER-Hoxb8 progenitors were differentiated and conditioned as in Figure 1A. Macrophages were subsequently co-cultured with apoptotic MDA-MB-231s that were labeled with ^13^C-Lysine to an incorporation rate > 99%. Naïve and efferocytotic macrophages from standard oxygen and physiological hypoxia conditions were harvested in the indicated oxygen environment prior to downstream processing and mass spectrometry analysis. Four independent experiments were analyzed for each condition. (Middle/Right) Based on our analyses of global proteomics data (see Figure S3), we observed differentially regulated mRNA transcripts (Core Programs; Middle) that were concomitantly differentially regulated at the protein level in apoptotic cell-naïve macrophages (Primed; Top Right) or differentially regulated mRNA transcripts that were not differentially regulated in apoptotic cell-naïve macrophages but instead were differentially regulated in efferocytotic macrophages (Poised; Bottom Right). Numbers represent individual genes/proteins differentially regulated, with light shading representing downregulated and dark shading representing upregulated. (B) <u>Representative genes/proteins of ‘primed’ and ‘poised’ states.</u> Shown are representative transcripts/proteins from each state observed in Figure 3A. PFKL/*Pfkl*, ATP-dependent 6-phosphofructokinase, liver type. SAMD9L/*Samd9l*, Sterile alpha motif domain-containing protein 9-like. Additional representative examples are presented in Figure S4. (C) <u>Temporal disruption of ‘poised’ state program translation impairs efferocytosis in chronic</u> <u>physiological hypoxia-conditioned macrophages.</u> (Left) Schematic illustrating the differential effect of cycloheximide (Chx) treatment on ‘primed’ versus ‘poised’ programs. (Right) Macrophages were differentiated and conditioned as in Figure 1A. Cycloheximide (or vehicle control) was added 1h prior to the start of an assays, then subsequently cultured with CypHer5E-labeled apoptotic Jurkat T cells at a 1:1 ratio for 1h in the presence of Chx or vehicle control. Efferocytosis was assessed via flow cytometry. Data represent three independent experiments. Data are shown as mean ± SEM. ***p < .001; ns = not significant. (D) <u>Time-lapse microscopy analysis of continual efferocytosis in chronic physiological hypoxia-conditioned macrophages with temporal disruption of ‘poised’ state program translation.</u> Experiments were performed as in Figure 1C. Shown are representative images (Left), quantification of the number of CypHer5E+ events per cell (Top Right), and analysis of the rate of apoptotic cell degradation (Bottom Right). For quantification, 70 efferocytotic macrophages from 5 vehicle-treated chronic physiological hypoxia scenes and 62 efferocytotic macrophages from 7 Chx-treated chronic physiological hypoxia scenes were analyzed. Data were binned as number of events per cell and presented as a fraction of 100%. For analysis of degradation rate, 38 (vehicle-treated) and 21 (Chx-treated) efferocytotic macrophages were analyzed. Time to degradation was defined as the time it takes to shrink an internalized corpse 50% after initial acidification (CypHer5E+). Data shown as mean ± SEM. ****p < .0001. (E) <u>Clearance of apoptotic thymocytes is perturbed *in vivo* in mice treated with cycloheximide.</u> Schematic of experimental design (Left) and summary plot (Right) of Annexin V+ 7-AAD+ (late/secondary apoptotic) cells from isolated thymocytes 6h post-dexamethasone (Dex) injection in vehicle (n = 5) or cycloheximide (Chx)-treated mice (n = 5). Data shown as mean ± SEM. ****p < .0001.

Our discovery of ‘poised’ state programs raised the question, what purpose does enhanced transcription of core functional genes without commensurate protein synthesis serve? One hypothesis is that translation of ‘poised’ state programs is necessary to support continuous uptake and degradation of apoptotic cells observed in chronic physiological hypoxia-conditioned macrophages. To test this, we took advantage of the properties of cycloheximide (Chx) to temporally disrupt translation with minimal effect on transcription (Figure 3C, left). Treatment of standard oxygen-conditioned macrophages with Chx immediately prior to culture with apoptotic cells did not affect efferocytosis (Figure 3C, right). Contrarily, treatment of chronic physiological hypoxia-conditioned macrophages with Chx immediately prior to culture with apoptotic cells resulted in a significant decrease in efferocytosis (Figure 3C). Furthermore, using time-lapse confocal microscopy, we tested if disrupting translation affects per cell engulfment and apoptotic cell digestion rate. Strikingly, we observed that chronic physiological hypoxia-conditioned macrophages with temporally disrupted translation engulfed significantly fewer apoptotic cells on a per cell basis (Chx-treated macrophages: ∼90% engulfed 2 or fewer corpses per cell compared to 50% of untreated macrophages which engulfed 3 or more corpses per cell; Figure 3D, top bar graph). Additionally, temporally disrupted translation resulted in a slower rate of apoptotic cell degradation (Figure 3D, bottom bar graph), suggesting that translation of the ‘poised’ program is essential for the rapid, continuous efferocytosis observed in chronic physiological hypoxia-conditioned macrophages.

Finally, we sought to determine if the ‘poised’ program is important for efferocytosis *in vivo*. Many of the tissues where macrophages perform high efferocytosis burden feature physiological hypoxia (Zago et al., 2021). For instance, macrophages in the thymus are responsible for clearing millions of apoptotic thymocytes daily (Breed et al., 2019; Stritesky et al., 2013) despite the thymus featuring physiological hypoxia (10mmHg, or ∼1%, O_2_ (Hale et al., 2002)). Taking advantage of its *in vivo* activity, we treated mice with Chx or vehicle prior to the induction of thymocyte apoptosis using dexamethasone (Figure 3E, left). Similar to our *in vitro* findings, mice treated with Chx exhibited significantly decreased clearance of apoptotic thymocytes, as indicated by increase in Annexin V+ 7AAD+ thymocytes (Figure 3E, right). Thus, the ‘poised’ program observed in chronic physiological hypoxia-conditioned macrophages is necessary for efficient continual efferocytosis *in vitro* and *in vivo*.

### Mapping the metabolic pathways unique to chronic hypoxia-conditioned macrophages

Given the highly coordinated regulation of specific metabolic pathways and processes in ‘primed’ macrophages, we further explored macrophage metabolism under chronic hypoxia. As part of our analysis of transcriptional programs in chronic physiological hypoxia-conditioned macrophages, we performed a network analysis of the differentially regulated metabolic programs in ‘primed’ macrophages (Figure 4A, Figure S4A). Our analysis revealed that the broader metabolic programs clustered into specific metabolic pathways and processes. For instance, carbohydrate metabolism clustered into three categories: upregulation of gluconeogenesis/ glucose transport, downregulation of one carbon/aldehyde metabolism, and mixed regulation of the pentose phosphate pathway, including upregulation of the oxidative stage and downregulation of the non-oxidative stage (PPP; Figure 4A, pathway highlighted in panel on the right). Consistent with our RNAseq and proteomics analyses, we observed significant differences in glucose uptake by macrophages conditioned in hypoxia (Figure 4B). However, contrary to our expectations, we observed (via enzymatic analysis) that chronic physiological hypoxia-conditioned macrophages consume significantly less glucose from media than macrophages in standard oxygen (Figure 4B, left graph). Furthermore, chronic physiological hypoxia-conditioned macrophages secrete less lactate (Figure 4B, right graph) and contain less ATP than macrophages under atmospheric (standard) oxygen (Figure 4C).

**Figure 4:**
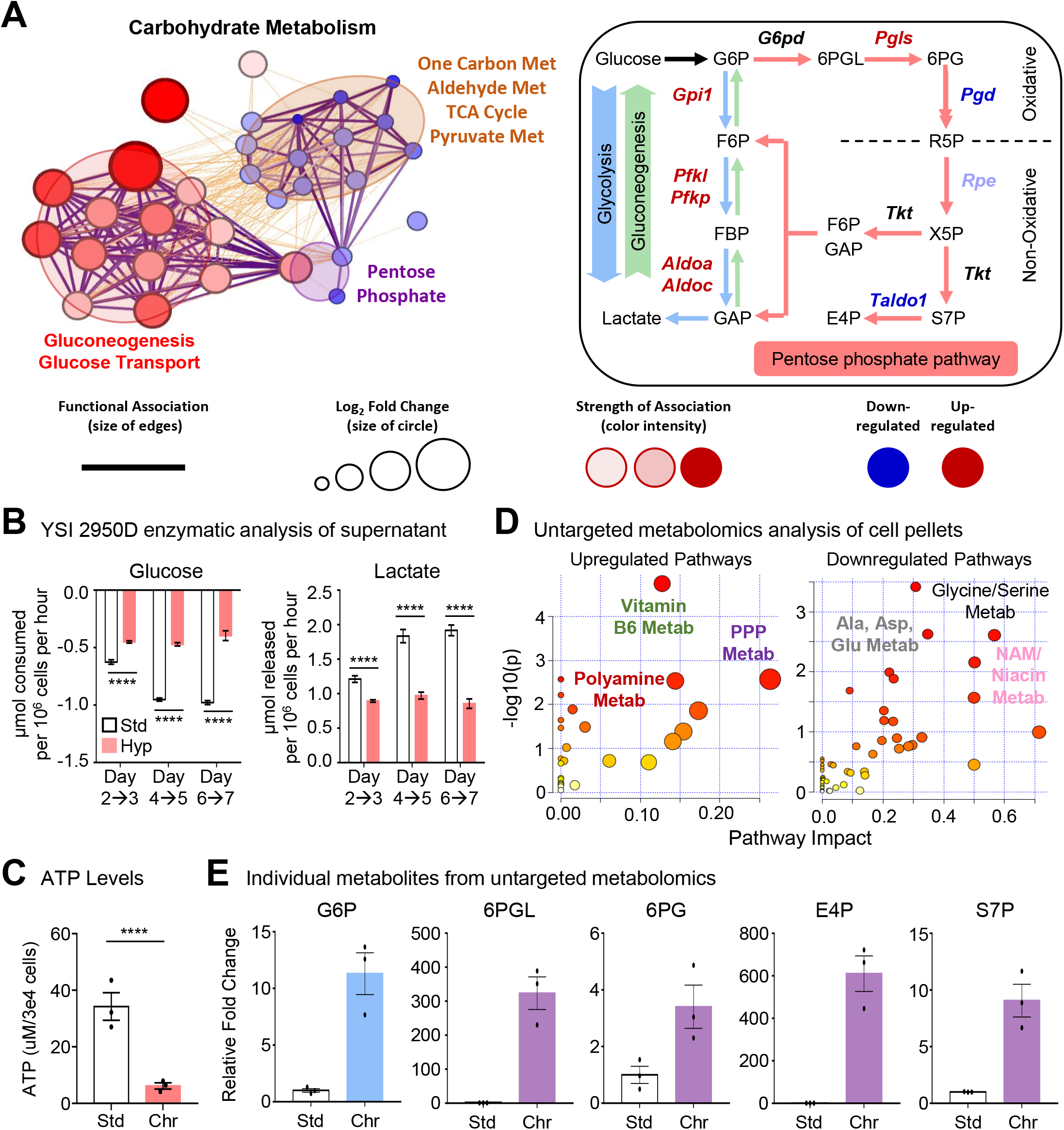
Metabolic pathway use in chronic physiological hypoxia-conditioned macrophages. (A) <u>Network analysis of differentially expressed metabolic transcripts identified in chronic</u> <u>physiological hypoxia-conditioned macrophages.</u> (Left) Metabolic genes that are differentially regulated in chronic physiological hypoxia-conditioned macrophages (Figure 2) are represented using network analysis to determine family clusters (shaded areas) and connectedness between individual genes. See also Figure S5A. (Right) Schematic of three main routes of glucose and glycolysis intermediate use: glycolysis (blue), gluconeogenesis (green), and pentose phosphate pathway (red). Genes in red are significantly upregulated whereas genes in blue are downregulated. (B) <u>Consumption of glucose and release of lactate by standard oxygen and physiological hypoxia-conditioned macrophages.</u> Macrophages were differentiated and cultured as in Figure 1A. Media was replaced on days 2, 4, and 6, and harvested 24h later (e.g., on days 3, 5, and 7). Media was analyzed using a 2950D Biochemistry Analyzer (YSI Life Sciences) for glucose and lactate levels compared relative to media controls conditioned in parallel. Two independent experiments were analyzed for each condition, with 4 biological replicates per experiment. Data are shown as mean ± SEM. ****p < .0001. (C) <u>Measurement of ATP levels in standard oxygen and physiological hypoxia-conditioned</u> <u>macrophages.</u> Macrophages were differentiated and cultured as in Figure 1A. Macrophages were collected on day 7 and ATP levels were analyzed. Three independent experiments were analyzed for each condition. Data are shown as mean ± SEM. ****p < .0001. (D) <u>Unique metabolomic profile in chronic hypoxia-conditioned macrophages.</u> Macrophages were differentiated and conditioned as in Figure 1A then isolated for untargeted metabolomic analysis. (Left) upregulated and (Right) downregulated pathways in chronic hypoxia-conditioned macrophages relative to standard oxygen and acute hypoxia-conditioned macrophages are shown (see Figure S4B for principal component analysis of all three conditions). Three independent experiments were performed for each condition. (E) <u>Representative individual metabolites from untargeted metabolomics analysis.</u> Shown are representative metabolites from significantly differentially regulated pathways from Figure 4C, specifically upper glycolysis (blue) and pentose phosphate pathway (PPP, purple). See also Figure S4C for additionally representative metabolites from upper glycolysis, PPP, lower glycolysis/TCA cycle (orange), and alanine/aspartate/glutamine metabolism (Ala, Asp, Glu Metabolism, gray). Data are shown as mean ± SEM. All metabolites are significant, p < .0001. G6P, glucose 6-phosphate. 6PGL, 6-phosphogluconolactone. 6PG, 6-phosphogluconate. E4P, erythrose 4-phosphate. S7P, sedoheptulose 7-phosphate.

Our findings suggesting that chronic physiological hypoxia-conditioned macrophages consume and utilize less glucose was surprising given previous studies of cellular metabolism under hypoxia (Denko, 2008; Sadiku and Walmsley, 2019; Taylor and Colgan, 2017). To further elucidate the metabolic state of chronic physiological hypoxia-conditioned macrophages, we performed untargeted metabolomics analysis of macrophages cultured in standard oxygen, conditioned in physiological hypoxia acutely, or conditioned chronically in physiological hypoxia. Similar to our RNAseq analysis, chronic physiological hypoxia-conditioned macrophages exhibited a different cellular metabolite profile than macrophages from either standard or acute hypoxia conditions (Figure S4B). Our analysis revealed significant upregulation of several pathways, including the pentose phosphate pathway (PPP), polyamine metabolism, and vitamin B6 metabolism (Figure 4D and 4E; Figure S4C). On the other hand, we observed downregulation of alanine/aspartate/glutamine metabolism, glycine/serine (one carbon) metabolism, and niacin/nicotinamide (vitamin B3) metabolism (Figure 4D and 4E; Figure S4C). We also observed decreased TCA cycle intermediates (Figure S4C), consistent with our observation that chronic physiological hypoxia-conditioned macrophages downregulate a mitochondrial metabolism transcriptional program (Figure S4A). Analysis of individual metabolites revealed relative enrichment of key PPP intermediates and the upstream glycolysis intermediates glucose 6-phosphate, fructose 6-phosphate, and fructose 1,6-bisphosphate but downregulation of/no change in downstream glycolysis intermediates, including glyceraldehyde 3-phosphate and pyruvate (Figure 4D; Figure S4C). Thus, chronic physiological hypoxia-conditioned macrophages induce or suppress distinct metabolic programs, in particular increased PPP activity despite decreased glucose uptake/lactate secretion and ATP synthesis.

### Macrophages adapt to chronic physiological hypoxia by co-opting a noncanonical pentose phosphate loop to ensure redox homeostasis

Beyond ATP generated during glycolysis, glucose also contributes to both redox homeostasis (e.g., NADPH generation) and nucleotide synthesis by shunting the glycolysis intermediate glucose 6-phosphate (G6P) into the pentose phosphate pathway (PPP; Figure 5A). Our finding that chronic physiological hypoxia-conditioned macrophages consume less glucose, secrete less lactate, contain less ATP, and accumulate fewer TCA cycle intermediates but significantly accumulate PPP intermediates suggests that macrophages may be utilizing glucose in an unconventional way. One possibility is that chronic physiological hypoxia-conditioned macrophages accumulate PPP intermediates because of more <u>efficient</u> use of glucose for PPP, i.e., more G6P enters the PPP per molecule of glucose. To test this hypothesis, we performed 1,2-^13^C-glucose tracing of standard oxygen- and chronic physiological hypoxia-conditioned macrophages. Isotope tracing analysis using 1,2-^13^C-glucose allowed us to determine how cells functionally use consumed glucose by measuring the fraction and relative abundance of metabolic intermediates downstream of glucose, including glycolysis, PPP, and the re-synthesis of glycolysis intermediates via gluconeogenesis (Figure 5A; (Jang et al., 2018)). In chronic physiological hypoxia-conditioned macrophages, we observed increased fractional enrichment and relative abundance of several intermediates that had cycled through the PPP (M+1 isotopologues, Figure 5A; Figure S5A), including fructose 6-phosphate (F6P), fructose 1,6-bisphosphate (FBP), and 3-phosphoglycerate (3PG). Additionally, chronic physiological hypoxia-conditioned macrophages exhibited increased relative abundance of the committed PPP intermediate sedoheptulose 7-phosphate (S7P; Figure S5A). Contrarily, we observed decreased fractional enrichment and relative abundance of the terminal glycolysis intermediates phosphoenolpyruvate (PEP) and lactate as well as decreased contribution of glucose to the glucose-alanine and TCA cycles in chronic physiological hypoxia-conditioned macrophages (Figure S5A). Importantly, and consistent with our LC-MS analysis of cell pellets, we detected significantly less M+1 (via PPP) and M+2 (via glycolysis) lactate in conditioned media from chronic physiological hypoxia-conditioned macrophages using carbon nuclear magnetic resonance (Figure 5B), suggesting that chronic physiological hypoxia-conditioned macrophages utilize less glucose for energy generation via glycolysis, instead <u>efficiently</u> shunting glucose into the PPP.

**Figure 5:**
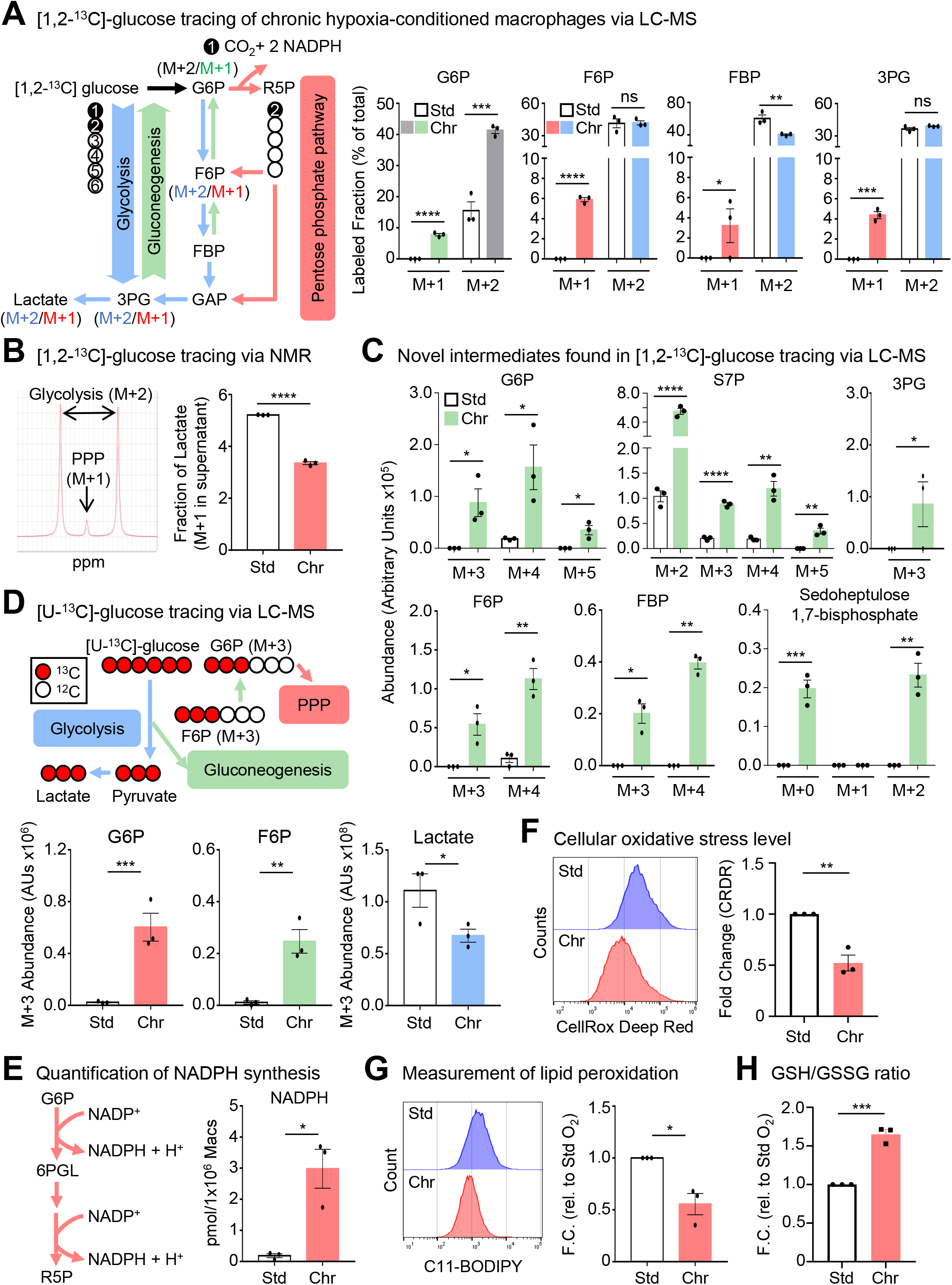
Macrophages co-opt noncanonical pentose phosphate loop for redox homeostasis. (A) <u>The pentose phosphate pathway is active in chronic physiological hypoxia-conditioned</u> <u>macrophages.</u> (Left) Schematic for isotopologue analysis of intermediates for pentose phosphate pathway (PPP; M+1) and glycolysis (M+2) in [1,2-^13^C]-glucose tracing experiments. (Right) Macrophages were differentiated and conditioned as in Figure 1A. On day 6, macrophages were cultured with media containing [1,2-^13^C]-glucose for 16h then analyzed via LC-MS. M+1 G6P, which arises via gluconeogenesis, is denoted in green. PPP intermediates (M+1) are denoted in red and glycolysis intermediates (M+2) are denoted in blue. Shown is the fractional enrichment of the indicated isotopologues. For relative abundance, see Figure S6A. Three independent experiments were performed for each condition. Data are shown as mean ± SEM. **p < .01; ***p < .001; ****p < .0001; ns = not significant. G6P, glucose 6-phosphate. F6P, fructose 6-phosphate. FBP, fructose 1,6-bisphosphate. 3PG, 3-phosphoglyceric acid. (B) <u>Lactate is not a major product of glucose metabolism in chronic physiological hypoxia-conditioned macrophages.</u> (Left) Shown are representative peaks for PPP (M+1)- and glycolysis (M+2)-derived lactate in media analyzed via NMR from experiments performed similar to Figure 5A. (Right) Cumulative data of the M+1 lactate fraction from our NMR analysis of macrophage-conditioned media relative to cell-free conditioned media. Data are from four biological replicates. Data are shown as mean ± SEM. ****p < .0001. (C) <u>Identification of flux through novel pentose phosphate pathway intermediates in chronic</u> <u>physiological hypoxia-conditioned macrophages.</u> Analysis of experiments performed in Figure 5A. Shown is summary data from our identification of isotopologues arising via gluconeogenesis (green). Additionally, we identified flux through the novel metabolite sedoheptulose 1,7-bisphosphate. Shown is the relative abundance of the indicated isotopologues. For fractional enrichment, see Figure S6B. Three independent experiments were analyzed for each condition. Data shown as mean ± SEM. *p < .05; **p < .01; ***p < .001; ****p < .0001. G6P, glucose 6-phosphate. S7P, sedoheptulose 7-phosphate. 3PG, 3-phosphoglyceric acid. F6P, fructose 6-phosphate. FBP, fructose 1,6-bisphosphate. (D) <u>Gluconeogenesis is prevalent in chronic physiological hypoxia-conditioned macrophages.</u> (Top) Schematic for [U-^13^C]-glucose flux analysis including routes through glycolysis, PPP, and gluconeogenesis. (Bottom) Experiments were performed identical to Figure 5A, except macrophages were incubated with [U-^13^C]-glucose for 16h then analyzed via LC-MS. M+3 G6P, which arises via PPP, is denoted in red. M+3 F6P, which arises via gluconeogenesis, is denoted in green. M+3 lactate, which arises via conventional glycolysis, is denoted in blue. Shown is the relative abundance of the indicated isotopologues. For fractional enrichment, see Figure S6C. Three independent experiments were performed for each condition. Data are shown as mean ± SEM. *p < .05; **p < .01; ***p < .001. G6P, glucose 6-phosphate. F6P, fructose 6-phosphate. (E) <u>NADPH synthesis is enhanced in chronic physiological hypoxia-conditioned macrophages.</u> (Left) Schematic of NADPH synthesis via pentose phosphate pathway. Macrophages were differentiated and conditioned as in Figure 5A. (Right) NADPH levels (pmol) were quantified relative to media-only controls. Three independent experiments were analyzed for each condition. Data shown as mean ± SEM. *p < .05; ****p < .0001. See also Figure S7A. (F) <u>Chronic physiological hypoxia-conditioned macrophages exhibit lower cellular oxidative stress</u> <u>compared to standard oxygen-conditioned macrophages.</u> Macrophages were differentiated and conditioned as in Figure 5A. Oxidative stress levels were measured by labeling macrophages with CellRox Deep Red and analysis via flow cytometry. Shown are representative FACS plots (Left) and summary plots of the MFI fold change relative to standard oxygen-conditioned macrophages (Right). Data are representative of three independent experiments per condition. Data are shown as mean ± SEM. **p < .01. (G) <u>Chronic hypoxia-conditioned macrophages exhibit lower lipid peroxidation compared to</u> <u>standard oxygen-conditioned macrophages.</u> Macrophages were differentiated and conditioned as in Figure 5A, incubated with C11-BODIPY 581/591 and analyzed via flow cytometry. Shown are representative FACS plots (Left) and fold change of lipid peroxidation level normalized to standard oxygen-conditioned macrophages. Data are representative of three independent experiment per condition. Data are presented as mean ± SEM. *p < .05. (H) <u>Analysis of the reduced/oxidized (GSH/GSSG) glutathione ratio in conditioned macrophages.</u> Macrophages were differentiated and conditioned as in Figure 5A and subsequently analyzed via plate reader for total glutathione, reduced glutathione (GSH), and oxidized glutathione (GSSG). Net relative luminescence unit (RLU) values were normalized to cell number. GSH/GSSG ratios were normalized to standard oxygen-conditioned macrophages. Data are from four independent experiments for each condition. Data are shown as mean ± SEM. ***p < .001. See also Figure S7E.

As part of our 1,2-^13^C-glucose tracing studies, we observed intermediates with ^13^C labeling beyond the first and second position (Figure 5C; Figure S5B). For instance, we observed increased fractional enrichment and relative abundance of M+3, M+4, and M+5 isotopologues of G6P in chronic physiological hypoxia-conditioned macrophages (Figure 5C; Figure S5B). The presence of these isotopologues could only arise if intermediates generated during glycolysis or PPP cycled back to G6P, through a process resembling gluconeogenesis in the liver (Hers and Hue, 1983). Unlike conventional gluconeogenesis, however, we did not detect newly synthesized glucose. Instead, together with G6P, we observe increased fractional enrichment and relative abundance of newly synthesized glycolytic intermediates F6P (M+3, M+4), FBP (M+3, M+4), and 3PG (M+3; Figure 5C). Additionally, we observed increased relative abundance and fractional enrichment of newly synthesized M+3, M+4, and M+5 isotopologues of S7P (Figure 5C; Figure S5B). In parallel, we performed uniformly-labeled (U)-^13^C-glucose tracing of standard oxygen- and chronic physiological hypoxia-conditioned macrophages to directly test glucose carbon recycling and turnover (Figure 5D). Similar to our 1,2-^13^C-glucose tracing studies, we observed less fractional enrichment and relative abundance of the glycolysis-generated M+3 lactate isotopologue in chronic physiological hypoxia conditioned macrophages (Figure 5D; Figure S5C). Furthermore, chronic physiological hypoxia-conditioned macrophages exhibited increased fractional enrichment and relative abundance of newly-synthesized M+3 G6P and F6P isotopologues (Figure 5D; Figure S5C). Of the additional possible fates for glucose, chronic physiological hypoxia-conditioned macrophages produced less alanine and decreased contribution to the TCA cycle, irrespective of route (via M+2 citrate or M+3 oxaloacetate, Figure S5D and S5E). Finally, glucose that enters the PPP can provide the five-carbon nucleoside sugar necessary for *de novo* nucleotide synthesis. We observed decreased *de novo* synthesis of both inosine monophosphate (IMP) and guanosine monophosphate (GMP; Figure S6F) in chronic physiological hypoxia-conditioned macrophages, suggesting that the adaptation macrophages undergo in response to physiological hypoxia includes a specific and efficient repurposing of glucose for the generation of NADPH.

Although the conventional PPP is thought to be important for inflammatory macrophage function, this often coincides with enhanced glycolysis, decreased G6P, and increased TCA cycle intermediates such as succinate and oxaloacetate (Artyomov and Van den Bossche, 2020; Makowski et al., 2020; Ryan and O’Neill, 2020). Contrarily, we observe increased PPP absent the other features present in inflammatory macrophages, suggesting that chronic physiological hypoxia-conditioned macrophages co-opt an unconventional PPP. In support of this, we detected M+0 and M+2 sedoheptulose 1,7-bisphosphate (SBP) only in chronic physiological hypoxia-conditioned macrophages (Figure 5C). SBP is an unusual metabolite, previously identified as a novel PPP intermediate in the liver (L-type PPP; (Cheng et al., 2019; Longenecker and Williams, 1980; Williams et al., 1985)). SBP is proposed to be synthesized via three different routes: 1) from dihydroxyacetone phosphate and erythrose 5-phosphate or 2) ribose 5-P via aldolase (e.g., ALDOA), and 3) from S7P via 6-phosphofructokinase (e.g., PFKL). We observed concomitant increases in mRNA and protein for aldolase (ALDOA and ALDOC) and 6-phosphofructokinase (PFKL and PFKP) (Figure 2). Interestingly, a recent report found that SBP accumulates in the hepatoma HepG2 line in response to oxidative stress (Cheng et al., 2019), suggesting that cells upregulate the noncanonical PPP to cope with contexts that pose a danger to redox homeostasis. Indeed, chronic physiological hypoxia-conditioned macrophages exhibited increased synthesis of NADPH and a decreased ratio of NADP+ to NADPH (Figure 5E, Figure S7A). Furthermore, macrophages exhibit lower cellular oxidative stress (Figure 5F) and lipid peroxidation (Figure 5G), further supporting the notion that macrophages adapt to physiological hypoxia by efficiently enhancing reductive potential and actively maintaining redox homeostasis.

Our RNAseq and proteomic analyses suggested that macrophages respond to chronic physiological hypoxia by regulating mitochondrial metabolism and mitochondrial biology (e.g., mitochondrial quality control) programs (Figure 2 and 3A; Figure S2, S3C, and S5A), which are major contributors to the redox state of a cell (Muri and Kopf, 2020). Subsequent analysis of chronic physiological hypoxia-conditioned macrophages revealed decreased mitochondrial mass and potential (Figure S7B and S7C), suggesting that macrophages limit mitochondrial processes that produce reactive species when exposed to prolonged physiological hypoxia. Additionally, chronic physiological hypoxia-conditioned macrophages exhibited decreased mitochondrial superoxide levels (Figure S6D). As further evidence of efficient metabolic adaptation to enhance reductive potential and maintain redox homeostasis, we observed an increased ratio of reduced glutathione (GSH) to oxidized glutathione (GSSG) (Figure 5H) despite decreased glutathione synthesis (Figure S7E) in chronic physiological hypoxia-conditioned macrophages relative to standard oxygen-conditioned macrophages. Collectively, our data indicate that macrophages adapt to chronic physiological hypoxia by preferentially shunting glucose into a noncanonical pentose phosphate loop that enhances reductive potential, including increased NADPH generation, to maintain low levels of cellular oxidative stress.

### The noncanonical pentose phosphate loop supports continual efferocytosis

We hypothesized that induction of ‘primed’ programs in macrophages exposed to prolonged physiological hypoxia directly support ‘poised’ functional programs, such as efferocytosis. To this end, we tested whether induction of noncanonical pentose phosphate pathway (PPP) activity, one of the chronic physiological hypoxia-induced ‘priming’ programs, directly affects efferocytosis. Specifically, we co-opted multiple strategies to target the PPP specifically *during* efferocytosis (Figure S8A). <u>First</u>, we used the well-characterized noncompetitive inhibitor of glucose-6-phosphate dehydrogenase (G6PD), G6PDi1, to block the NADPH-producing conversion of G6P to 6-phosphogluconolactone (6PGL) which resulted in decreased efferocytosis by chronic physiological hypoxia-conditioned macrophages but not macrophages cultured in standard oxygen (Figure S8B, left bar graph). <u>Second</u>, we utilized a second small molecule inhibitor, 6-aminonicotinamide (6-AN), to target G6PD activity. Similar to G6PDi-1, 6-AN-mediated inhibition of G6PD resulted in significantly decreased efferocytosis in chronic physiological hypoxia-conditioned macrophages (Figure S8B, center bar graph). <u>Third</u>, to test the contribution of gluconeogenesis to efferocytosis, we targeted fructose 1,6-bisphosphatase-1 (FBP1) activity using the selective inhibitor FBPi, which blocks the rate-limiting conversion of fructose 1,6-bisphosphate (FBP) to fructose 6-phosphate (F6P). Consistent with our tracing studies suggesting that PPP intermediates cycle back to G6P, we found that FBP1 inhibition led to decreased efferocytosis exclusively in chronic physiological hypoxia-conditioned macrophages (Figure S8B, right bar graph). Collectively, our combination of approaches temporally targeting components of the noncanonical PPP loop suggest that macrophages depend on this loop for efficient efferocytosis under prolonged physiological hypoxia.

Considering our previous observation that chronic physiological hypoxia-conditioned macrophages exhibit enhanced continual efferocytosis, we queried whether PPP activity also supports continual efferocytosis. Temporal targeting of G6PD activity not only suppressed engulfment of the first corpse but also reduced engulfment of subsequent corpses (Figure 6A), suggesting that PPP activity is required for continual efferocytosis by chronic physiological hypoxia-conditioned macrophages. Continual efferocytosis depends on a phagocytes ability to rapidly mature the phagolysosome and degrade apoptotic cells, as observed in chronic physiological hypoxia-conditioned macrophages (Figure 1E and 1F). We subsequently tested whether temporal perturbation of the PPP affects per cell engulfment and rate of apoptotic cell digestion. Time-lapse confocal microscopy revealed three key findings. <u>First</u>, inhibition of G6PD activity resulted in fewer macrophages engulfing apoptotic cells, consistent with our population-level analysis of efferocytosis (Figure 6B). <u>Second</u>, we observed that macrophages engulf significantly fewer apoptotic cells on a per cell basis when G6PD activity was inhibited (6-AN-treated macrophages: ∼80% engulfed 2 or fewer corpses per cell compared to 70% of untreated macrophages which engulfed 3 or more corpses per cell) (Figure 6B, left bar graph). <u>Third</u>, inhibition of G6PD activity resulted in a slower rate of apoptotic cell degradation (Figure 6B, right bar graph), essentially reversing the ‘priming’ adaptations macrophages make under prolonged physiological hypoxia. Finally, we sought to determine if temporal perturbation of the ‘primed’ program affects efferocytosis *in vivo*. To this end, we treated mice with 6-AN or vehicle prior to the induction of thymocyte apoptosis using dexamethasone (Figure 6C, left). Similar to our *in vitro* findings, mice treated with 6-AN exhibited significantly decreased clearance of apoptotic thymocytes, as indicated by increase in Annexin V+ 7AAD+ thymocytes (Figure 6C, right).

**Figure 6:**
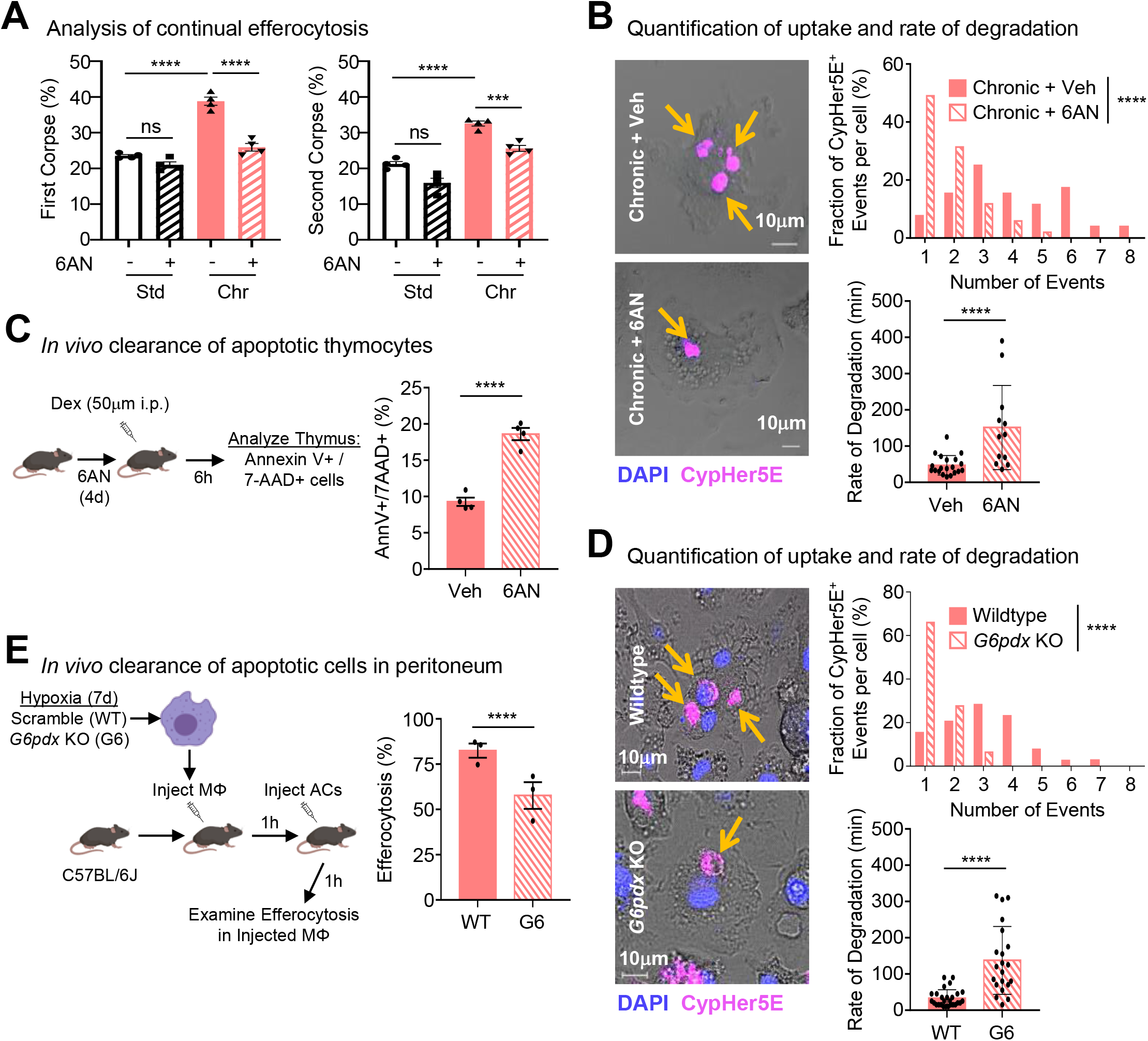
Noncanonical pentose phosphate loop supports continual efferocytosis. (A) <u>Perturbing PPP metabolism impairs continual efferocytosis in chronic physiological hypoxia-conditioned macrophages.</u> Macrophages were differentiated and conditioned as in Figure 1A then subsequently cultured with CFSE-labeled apoptotic MDA-MB-231 cells at a 1:1 ratio for 1h. Uncleared apoptotic cells were removed, and macrophages were either analyzed via flow cytometry (First Corpse, Left) or rested for 1.5h. Rested macrophages were then cultured with CypHer5E-labeled apoptotic MDA-MB-231 cells at a 1:1 ratio for 1h and analyzed via flow cytometry (Second Corpse, Right). Data represent three independent experiments. Data are shown as mean ± SEM. ***p < .001; ****p < .0001; ns = not significant. (B) <u>Time-lapse microscopy analysis of continual efferocytosis in chronic physiological hypoxia-conditioned macrophages with perturbed PPP metabolism.</u> Experiments were performed as in Figure 1E and 1F with the inclusion of 6AN and vehicle control treatment conditions. Shown are representative images (Left), quantification of the number of CypHer5E+ events (Top Right), and analysis of the rate of apoptotic cell degradation (Bottom Right). For quantification, 53 efferocytotic macrophages from 5 vehicle-treated chronic hypoxia scenes and 51 efferocytotic macrophages from 7 6AN-treated chronic hypoxia scenes were analyzed. Data were binned as number of events per cell and presented as a fraction of 100%. For analysis of degradation rate, 21 (vehicle-treated) and 13 (6AN-treated) efferocytotic macrophages were analyzed. Time to degradation was defined as the time it takes to shrink an internalized corpse 50% after initial acidification (CypHer5E+). Data shown as mean ± SEM. ****p < .0001. (C) <u>Clearance of apoptotic thymocytes is decreased *in vivo* in mice with perturbed PPP</u> <u>metabolism.</u> Schematic of experimental design (Left) and summary plot (Right) of Annexin V+ 7-AAD+ (late/secondary apoptotic) cells from isolated thymocytes 6h post-dexamethasone (Dex) injection in vehicle (n = 4) or 6AN-treated mice (n = 4). Data shown as mean ± SEM. ****p < .0001. (D) <u>Time-lapse microscopy analysis of continual efferocytosis in chronic physiological hypoxia-conditioned macrophages with genetic targeting of *G6pdx*.</u> Experiments were performed as in Figure 6B but with Cas9+ Hoxb8 macrophages bearing scramble (WT) or G6pdx (G6) small guides. Shown are representative images (Left), quantification of the number of CypHer5E+ events (Top Right), and analysis of the rate of apoptotic cell degradation (Bottom Right). For quantification, 39 efferocytotic macrophages from 5 WT chronic physiological hypoxia scenes and 47 efferocytotic macrophages from 7 G6 chronic physiological hypoxia scenes were analyzed. Data were binned as number of events per cell and presented as a fraction of 100%. For analysis of degradation rate, 27 (WT) and 21 (G6) efferocytotic macrophages were analyzed. Time to degradation was defined as the time it takes to shrink an internalized corpse 50% after initial acidification (CypHer5E+). Data shown as mean ± SEM. ****p < .0001. (E) <u>Clearance of apoptotic T cells is decreased in macrophages with genetic disruption of *G6pdx*</u> *<u>in vivo</u>*. (Left) Schematic of experimental design. Mice were first i.p. injected with Cas9/GFP+ Hoxb8 macrophages bearing scramble (WT) or *G6pdx* (G6) small guides 1h prior to i.p. injection of CypHer5E-labeled apoptotic cells (ACs). After 1h, peritoneal GFP+ CD11b+ F4/80+ macrophages were analyzed for efferocytosis. Shown is a summary plot of the rate of efferocytosis by GFP+ CD11b+ F4/80+ macrophages in WT mice (n=3) and G6 mice (n=3). ****p < .0001. Data shown as mean ± SEM. ****p < .0001.

An essential step in the oxidative PPP is the conversion of G6P into 6PGL mediated by the enzyme G6PD (Figure S8A). We next sought to determine if genetic targeting of *G6pdx*, the mouse isoform of G6PD, affects efferocytosis by chronic physiological hypoxia-conditioned macrophages. Similar to our results with the small molecule 6-AN, targeting of *G6pdx* (Figure S8C) revealed several striking findings. <u>First</u>, macrophages lacking *G6pdx* engulfed significantly fewer apoptotic cells on a per cell basis (*G6pdx*-deficient macrophages: ∼90% engulfed 2 or fewer corpses per cell compared to 65% of scramble control macrophages which engulfed 3 or more corpses per cell) (Figure 6D, top graph). <u>Second</u>, macrophages lacking *G6pdx* degraded apoptotic cells at a slower rate (Figure 6D, bottom graph), essentially reversing the ‘priming’ adaptations macrophages make under prolonged physiological hypoxia. <u>Finally</u>, *G6pdx*-deficient macrophages engulfed significantly fewer apoptotic cells than control macrophages *in vivo* (Figure 6E). Thus, the ‘primed’ program, in particular the noncanonical pentose phosphate loop, observed in chronic physiological hypoxia-conditioned macrophages is also necessary for efficient continual efferocytosis *in vitro* and *in vivo*.

### Noncanonical pentose phosphate loop supports phagolysosomal maturation and prevents redox crisis in efferocytotic macrophages

Phagolysosomal maturation and subsequent apoptotic cell degradation requires NADPH oxidase activity (Bagaitkar et al., 2018) but if not buffered against, can lead to redox crisis featuring excessive/runaway lysosomal acidification and oxidative stress (Mantegazza et al., 2008; YvanCharvet et al., 2010). NADPH, therefore, serves as both a substrate for reactive oxygen species production and an essential reducer of antioxidant molecules. NADPH is produced through three main pathways: the conversion of malate to pyruvate via malic enzyme 1 (ME1) activity, through conversion of isocitrate into alpha-ketoglutarate via isocitrate dehydrogenase 1 (IDH1) activity, and during the oxidative phase of the PPP. Macrophages under atmospheric (standard) oxygen conditions produce minimal NADPH (Figure 7A; (Yvan-Charvet et al., 2010)) and possibly utilize one of the oxygen-dependent TCA cycle-related routes (ME1 or IDH1) to generate NADPH during efferocytosis *in vitro* as well as by macrophages localized to tissues with higher oxygen availability (Zhang et al., 2019). Consistent with our finding that chronic physiological hypoxia-conditioned macrophages exhibit decreased glucose flux into the TCA cycle (Figure S6A and S6E) and increased flux into the PPP (Figure 5A and 5C), we observed concomitant PPP-dependent production of NADPH (Figure 7A). Although NADPH production remained abundant during efferocytosis by chronic physiological hypoxia-conditioned macrophages (Figure 7B, compare solid red and purple bars), NADPH pools were further depleted in efferocytotic macrophages with temporally disrupted 6PDG activity (Figure 7B, compare solid and striped purple bars) and decreased *G6pdx* levels (Figure 7C). Thus, chronic physiological hypoxia-conditioned macrophages boost PPP-dependent NADPH production prior to efferocytosis which remains active during efferocytosis.

**Figure 7:**
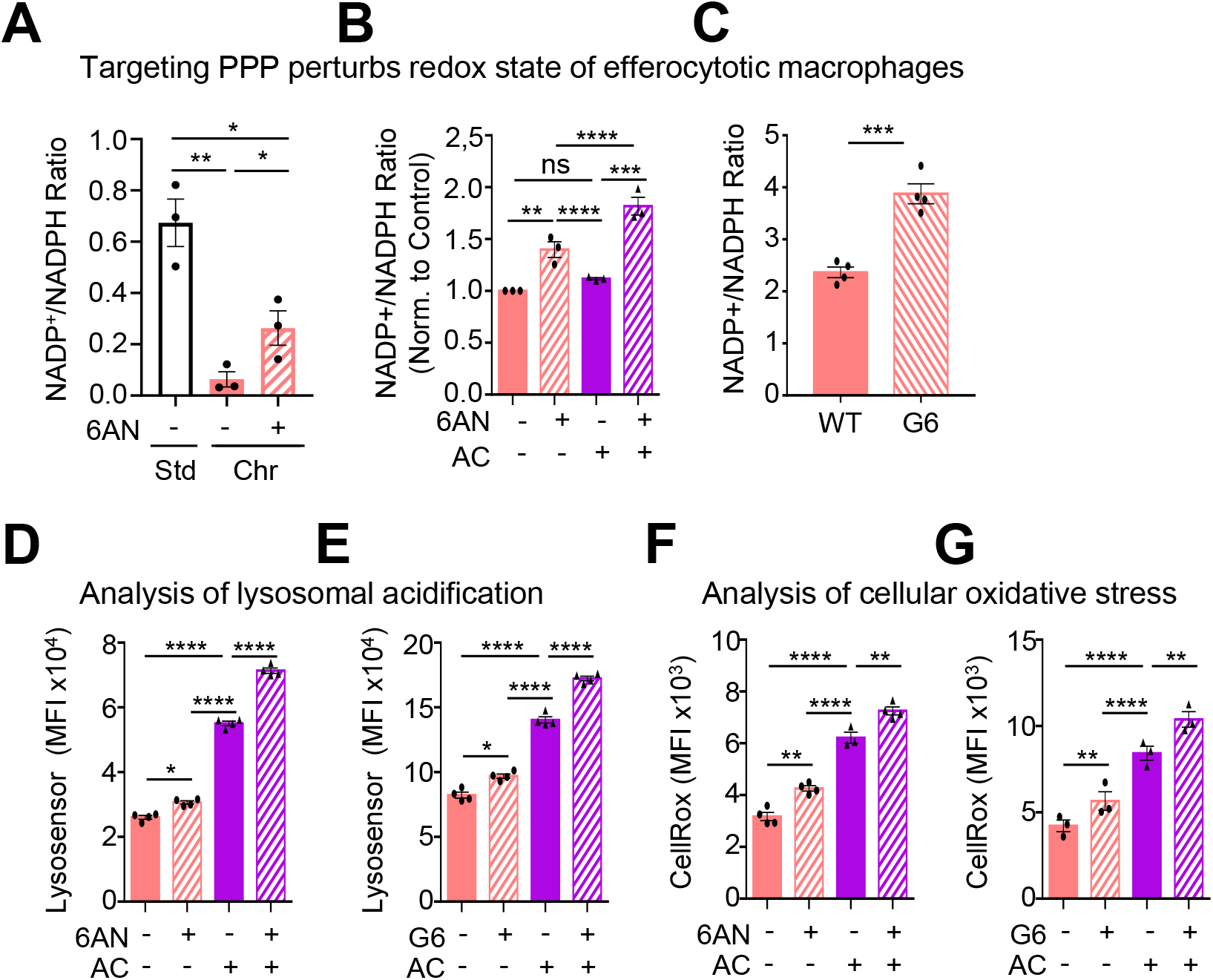
Noncanonical pentose phosphate loop prevents redox crisis in efferocytotic macrophages. (A) <u>Temporal perturbation of PPP metabolism reduces NADPH production in chronic hypoxia-conditioned macrophages.</u> Macrophages were differentiated and conditioned as in Figure 1A. Indicated treatments (or vehicle controls) were added prior to the start of an assays. NADP^+^ and NADPH relative luminescence units were detected using a plate reader. Data are from three independent experiments. Data are plotted as the ratio of NADP^+^ to NADPH and presented as mean ± SEM. *p < .05; **p < .01. (B) <u>Chronic physiological hypoxia-conditioned macrophages consume PPP-dependent NADPH</u> <u>during efferocytosis.</u> Macrophages were differentiated and conditioned as in Figure 7A. Conditioned macrophages were cultured with apoptotic MDA-MB-231 cells at a 1:1 ratio for 1h. Uncleared apoptotic cells were removed, and macrophages were lysed using a basic solution (0.2N NaOH, 1% DTAB) in a hypoxia chamber set to 1% O_2_. NADP^+^ and NADPH relative luminescence units were detected using a plate reader. Data are from three independent experiments. Data are plotted as the ratio of NADP^+^ to NADPH and presented as mean ± SEM. **p < .01; ***p < .001; ****p < .0001. ns = not significant. (C) <u>Genetic targeting of *G6pdx* reduces NADPH production in chronic hypoxia-conditioned</u> <u>macrophages.</u> Macrophages were differentiated and conditioned as in Figure 7A except using Cas9/GFP+ Hoxb8 macrophages bearing scramble (WT) or *G6pdx* (G6) small guides. NADP^+^ and NADPH relative luminescence units were detected using a plate reader. Data are from three independent experiments. Data are plotted as the ratio of NADP^+^ to NADPH and presented as mean ± SEM. ***p < .001. (D, E) <u>Lysosomal acidity increases in chronic physiological hypoxia-conditioned efferocytotic</u> <u>macrophages with temporally-perturbed PPP and genetically-targeted *G6pdx*.</u> Macrophages were differentiated and conditioned as in Figure 7A using 6AN (or vehicle)-treated macrophages or Cas9/GFP+ Hoxb8 macrophages bearing scramble (WT) or *G6pdx* (G6) small guides (**E**). Conditioned macrophages were cultured with CypHer5E^+^-labeled apoptotic MDA-MB-231 cells at a 1:1 ratio for 1h. Uncleared apoptotic cells were removed, and macrophages were stained with Lysosensor Green DND-189. CypHer5E^+^ macrophages were gated as actively efferocytotic for analysis of lysosomal acidification. Shown are representative FACS plots (Left) and MFI (Right) from three independent experiments per condition. Data are shown as mean ± SEM. *p < .05; ****p < .0001. (F, G) <u>Cellular ROS increases in chronic physiological hypoxia-conditioned efferocytotic</u> <u>macrophages with temporally-perturbed PPP and genetically-targeted *G6pdx*.</u> Macrophages were differentiated and conditioned as in Figure 7A using 6AN (or vehicle)-treated macrophages (**F**) or Cas9/GFP+ Hoxb8 macrophages bearing scramble (WT) or *G6pdx* (G6) small guides (**G**). Conditioned macrophages were cultured with CypHer5E^+^-labeled apoptotic MDA-MB-231 cells at a 1:1 ratio for 1h. Uncleared apoptotic cells were removed, and macrophages were stained with CellRox Deep Red. CypHer5E^+^ macrophages were gated as actively efferocytotic for analysis of cellular ROS. Shown are representative FACS plots (Left) and MFI (Right) from three independent experiments per condition. Data are shown as mean ± SEM. **p < .01; ****p < .0001.

Our observation that chronic physiological hypoxia-conditioned macrophages increase PPP-dependent NADPH production both prior to and during efferocytosis raises the hypothesis that the noncanonical PPP loop, a core ‘primed’ program, directly supports efferocytosis, a core ‘poised’ program. To test this hypothesis, we sought to determine if temporal perturbation or genetic disruption of the PPP also leads to excessive/runaway lysosomal acidification and oxidative stress in efferocytotic macrophages. In apoptotic cell-naïve macrophages, inhibition of 6PDG activity resulted in a modest, albeit significant, increase in lysosomal acidity (Figure 7D; Figure S8D, compare red bars). On the other hand, efferocytotic macrophages with perturbed 6PDG activity displayed significantly higher lysosomal acidity than untreated apoptotic cell-naïve and efferocytotic macrophages (Figure 7D; Figure S8D, purple striped bar). We observed similar results in chronic physiological hypoxia-conditioned macrophages with genetically-disrupted of *G6pdx* (Figure 7E). Our observation that disruption of PPP activity alters basal and, to a greater extent, efferocytotic lysosomal acidity is consistent with previous work demonstrating that pharmacological targeting of NADPH oxidase activity results in increased lysosomal acidity in both resting and efferocytotic macrophages (Mantegazza et al., 2008). Finally, we tested the hypothesis that disrupted PPP activity leads to increased oxidative stress in efferocytotic macrophages. Similar to our finding of increased lysosomal acidity, we observed that temporal perturbation of 6PDG activity resulted in significantly increased lipid peroxidation (Figure S8E, purple striped bar) and cellular oxidative stress (Figure 7F; Figure S8F, purple striped bar) in chronic physiological hypoxia-conditioned efferocytotic macrophages. Importantly, genetic disruption of G6pdx also resulted in higher oxidative stress (Figure 7G) in chronic physiological hypoxia-conditioned efferocytotic macrophages. Thus, our data suggest that PPP-mediated NADPH production, a core ‘primed’ program, supports enhanced efferocytosis, a core ‘poised’ program, by providing the NADPH necessary for both rapid phagolysosomal maturation and protection from excessive lysosomal acidification and oxidative stress under settings of prolonged physiological hypoxia.

## Discussion

Professional phagocytes, such as macrophages, reside in tissues throughout the body in limited nutrient environments often for long periods of time (Blériot et al., 2020; Guilliams et al., 2020). This is especially true for oxygen, which ranges between 38 mmHg (∼5%) and 10 mmHg (∼1%, physiological hypoxia; (Carreau et al., 2011)). Despite the relative dearth of oxygen, phagocytes remain responsible for removing millions of apoptotic cells daily (Norris et al., 2019), a process that is metabolically demanding and potentially perilous (Trzeciak et al., 2021). How phagocytes adapt to continuous inhabitance in physiological hypoxic environments and how such adaptations inform clearance of apoptotic cells (‘efferocytosis’) remains unexplored. Here, we made the striking observation that macrophages conditioned in prolonged physiological hypoxia (∼1% O_2_) *in vitro* or *in vivo* are better phagocytes, both engulfing more apoptotic cells and degrading internalized apoptotic cells faster. As part of an adaptive response to prolonged physiological hypoxia, macrophages induce two distinct states that support efficient efferocytosis. The first state, which we term ‘primed’, is characterized by concomitant changes in mRNA and protein programs, especially metabolic programs, in apoptotic cell-naïve macrophages that remain induced during efferocytosis. The second state, which we term ‘poised’, is characterized by transcription, but not translation, of phagocyte function programs (e.g., phagocytosis programs) in apoptotic cell-naïve macrophages that are subsequently translated during efferocytosis. We found that both primed and poised state programs support enhanced continual efferocytosis by macrophages residing under chronic physiological hypoxia. Importantly, we found that this ‘primed’ state, in particular induction of a noncanonical pentose phosphate pathway loop that generates abundant NADPH, directly supports enhanced, continual efferocytosis (a core ‘poised’ state program). Based on our findings, we propose that phagocytes that reside in physiological hypoxic environments, such as highly phagocytic macrophages in the bone marrow or spleen (Norris et al., 2019; Sender and Milo, 2021), adopt distinct states that both ensure their own fitness and their readiness to perform core homeostatic functions.

Previous analysis of phagocytosis in different oxygen levels has produced mixed findings. For instance, past studies have suggested that acute hypoxia enhances engulfment of opsonized targets (Acosta-Iborra et al., 2009), bacteria (Anand et al., 2007; Fritzenwanger et al., 2011), and apoptotic cells (Norris et al., 2019), whereas other studies have suggested that acute hypoxia has no effect on phagocytosis (Dehn et al., 2016; Fritzenwanger et al., 2011; Lin et al., 2018) or decreases clearance of apoptotic cells (Marsch et al., 2014), and that macrophages in hypoxic regions may produce less of the anti-inflammatory mediatory IL-10 typically produced during efferocytosis (Zhang et al., 2019). Much of the contradiction might relate to variations in how hypoxia was induced/maintain or for how long cells were cultured in hypoxia, typically ranging from 3h to 24h. Indeed, significant cellular changes can occur, including degradation of HIF proteins and HIF-dependent transcriptional programs, as quickly as 24h after introduction to hypoxia (Ginouvès et al., 2008; Jain et al., 2020). Although we observed a modest increase in population-level efferocytosis in macrophages exposed to acute physiological hypoxia, these cells did not exhibit enhanced lysosomal degradation. Instead, the most dramatic differences in both uptake and degradation of apoptotic cells were observed in macrophages conditioned in chronic physiological hypoxia. It will be interesting to further interrogate the importance of prolonged physiological hypoxia, including identified ‘primed’ and ‘poised’ programs, especially considering phagocytes residing in physiologically hypoxic tissues are often responsible for clearing biological material beyond conventional apoptotic cells (Trzeciak et al., 2021; Zago et al., 2021).

There is increased interest in how cells adapt metabolically to chronic hypoxia. For instance, Mootha and colleagues found that transformed cells (e.g., K562 cells) predominantly depend on lipid synthesis-and peroxisome biology-associated genes but not mitochondrial or Fe-S biosynthesis genes for survival in chronic hypoxia (Jain et al., 2020). In the present study, we observed downregulation of transcripts and proteins associated with mitochondrial and lipid metabolism as well as alterations in oxidative state, mitochondrial content, and membrane potential, similar to changes observed in chronic hypoxia-conditioned THP1 monocyte-derived macrophages (Fuhrmann et al., 2013). Unique in the current study, we observed that chronic physiological hypoxia-conditioned macrophages upregulate transcripts and proteins involved in glucose metabolism/gluconeogenesis as well as shunted glucose into a noncanonical pentose phosphate pathway (PPP) loop, supporting the hypothesis that cells residing in physiological hypoxia adapt by dramatically altering their subcellular biology to favor mitochondria-independent metabolic processes.

Despite the growing body of work detailing how cells undergo metabolic adaptation to chronic hypoxia, understanding how such changes inform cellular function remains largely unexplored. Importantly, we move beyond a phenomenological understanding of cellular metabolic adaption to chronic physiological hypoxia by mechanistically linking the changes in macrophage metabolism to enhanced apoptotic cell uptake and degradation. Unexpectedly, we found that macrophages under prolonged physiological hypoxia limit glucose flux to most major pathways including anaerobic glycolysis and the TCA cycle. Instead, macrophages shunt glucose into a noncanonical PPP loop involving repeated cycling of PPP intermediates back to G6P via a process resembling gluconeogenesis. Importantly, we found that this PPP loop is necessary for generation of NADPH in apoptotic cell-naïve macrophages and appears to provide the NADPH necessary for appropriate phagolysosomal maturation and apoptotic cell degradation. Collectively, we speculate that the adaptations observed in our study represent normal physiology of tissue-resident macrophages residing in physiological hypoxic tissues.

Previous studies of neutrophils and LPS-stimulated macrophages, both of which are thought to exist in an inflammatory state, have been shown to rely on PPP for function under atmospheric (standard) oxygen (Baardman et al., 2018; Gonçalves et al., 2020; O’Neill et al., 2016). Although this raises the question of whether macrophages conditioned in chronic physiological hypoxia are also inflammatory, several pieces of evidence counter this notion. First, it was previously shown that inflammatory macrophages, such as those elicited by LPS, have decreased efferocytosis potential compared to apoptotic cell-naïve macrophages (Feng et al., 2011; McPhillips et al., 2007; Michlewska et al., 2009; Thorp et al., 2011) whereas we observe increased efferocytosis potential in chronic physiological hypoxia-conditioned macrophages. Second, the transcriptional and translational programs of chronic physiological hypoxia-conditioned macrophages are similar to those observed in canonical pro-resolving macrophages, including expression of wound healing and anti-inflammatory programs highlighted by robust induction of Arginase 1 (ARG1). Finally, inflammatory macrophages and neutrophils exhibit increased glucose uptake, increased aerobic glycolysis and glucose-derived pyruvate generation, and increased TCA cycle activity (Cameron et al., 2019; Di Gioia et al., 2020; Liu et al., 2016), whereas chronic physiological hypoxia-conditioned macrophages display none of these changes. Although it remains possible that chronic physiological hypoxia-conditioned macrophages exhibit some features of an inflammatory response, our findings suggest that the transcriptional/translational programs and metabolic pathways induced under prolonged physiological hypoxia are distinct from those observed in canonical pro-inflammatory macrophages.

A series of important questions arise from our work related to our observation that chronic physiological hypoxia seems to induce both ‘primed’ and ‘poised’ states in macrophages. For instance, what macrophage functions, beyond efferocytosis, are ‘poised’ under chronic physiological hypoxia? Efferocytosis is generally a core macrophage function shared across tissue-resident macrophage populations, but are there functions that are informed by a combination of oxygen availability and tissue-specific factors (e.g., presence of heme)? How do changes in oxygen availability during development inform efferocytosis potential? Finally, how are these poised programs regulated, for example, are they maintained by ribosomal stalling or increased chromatin accessibility/transcription? Although beyond the scope of the work presented here, answers to these questions could provide important clues on how macrophage function in specific environments to maintain tissue homeostasis.

## STAR Methods

### Contact for Reagent and Resource Sharing

For further information and requests for resources and reagents should be directed to the Lead Contact, Justin S. A. Perry.

### Standard Oxygen and Hypoxia Conditioning

To model standard oxygen, acute hypoxia, and chronic hypoxia environments, we optimized protocols using the BioSpherix Xvivo X3 system, which allows us to dynamically regulate CO_2_ and O_2_ (down to 0.1%) in four different chambers without exposing cells to atmosphere. Primary macrophage progenitors immortalized with a modified estrogen receptorHoxb8 fusion (ER-Hoxb8) were generated as previously reported (Wang et al., 2006) from C57BL/6J mice. Progenitors were maintained in RPMI 1640 containing 5% heat-inactivated fetal bovine serum, 5% P885L (GM-CSF producing)-conditioned media, and 0.5μM β-estradiol (Sigma) in standard oxygen (∼21% O_2_). To differentiate into primary macrophages, progenitors were first washed three times with cold PBS to remove β-estradiol. Progenitors were then cultured in a-MEM containing 5% heat-inactivated fetal bovine serum, 1% PenicillinStreptomycin-Glutamine (100X), and 10% L929 (M-CSF producing)-conditioned media. On day 6 of differentiation, mature macrophages were replated in non-TC treated plates and cultured in either standard oxygen or hypoxia (1% O_2_) for 7 days prior to use in efferocytosis assays. For some experiments, progenitors were instead cultured in standard oxygen or hypoxia for 7 days prior to differentiation and use in efferocytosis assays. In all cases, media was replenished every other day and cells were incubated @37°C.

### Induction of Apoptosis

For thymocyte engulfment, thymi from 6-week-old mice were harvested and crushed between frosted slides to release thymocytes. Cells were then treated with dexamethasone (50 M) for 4 hours prior to downstream use. Jurkat T lymphoma, MDA-MB-231, BrM2, and E0771 cells, were induced to undergo apoptosis by treatment with 150mJ cm^−2^ (Jurkat cells) or 650mJ cm^−2^ ultraviolet C irradiation (Stratalinker) at ∼5×10^6^ cell density per 10cm non-TC treated dish. Then, cells were incubated for 4h, 8h, 16h, or 24h before downstream use. Apoptosis was confirmed via Annexin V/7-AAD staining to be greater than 70% Annexin V+ single-positive (Figure S1A).

### *In Vitro* Efferocytosis Assays

Apoptotic cells were stained with either 1μM CypHer5E (Cytiva) or 50μM TAMRA-SE (ThermoFisher) in serum-free HBSS for 45min. Then, cells were washed by incubating in serumcontaining assay media for an additional 25min. Apoptotic cells were co-cultured with macrophages at a 1:1 phagocyte to target ratio. All efferocytosis assays included four technical replicates for each condition with at least three biological replicates. For all pharmacological studies, phagocytes were pre-incubated with the compounds listed below or vehicle (DMSO) for 16h or 6d prior to addition of apoptotic cells. Macrophages were treated with the following: Cycloheximide (Chx; Sigma, 100μM), 6-AN (MedChemExpress, 100μM), G6PDi-1 (Cayman Chemical, 50μM), or FBP1i (Cayman Chemical, 25μM). Macrophages were examined via microscopy to ensure no gross morphological changes or cell death had occurred due to drug treatment. Macrophages and apoptotic cells were co-cultured for 1h. Apoptotic cells were subsequently removed via three washes with cold assay media and phagocytes were harvested using a cell scraper (Biotium) and assessed by flow cytometry. For continual efferocytosis, the first round of apoptotic cells was labeled with CFSE (ThermoFisher) and co-cultured with macrophages at a 1:1 ratio for 1h. Then, apoptotic cells were washed away using cold assay media. Macrophages were subsequently rested for 1.5h prior to co-culture with a second round of apoptotic cells that were labeled with CypHer5E. Continual efferocytosis was assessed by FACS as CFSE+ (1st corpse uptake) and CFSE+ CypHer5E+ (2nd corpse uptake) macrophages. FACS analysis was performed using an Attune NxT Flow Cytometer (ThermoFisher) and analyzed using FlowJo 10.7.1.

### Time-lapse Confocal Microscopy

#### Equipment

Experiments to evaluate per cell uptake and degradation were performed on a Zeiss Axio Observer.Z1 7 inverted fluorescence microscope equipped with a Zeiss 20x PlanApo (0.8 NA) objective, a 6-channel, 7-laser LSM 980 (405nm, 445nm, 488nm, 514nm, 561nm, 594nm, 639nm), and a Airyscan 2 multiplex detector. The imaging stage is fully encased in a black-out environmental chamber and the Z PIEZO stage is fully encapsulated with a heating/gas-controlled insert, together allowing us to control the temperature, humidity, CO_2_, and O_2_. Time-lapse experiments were acquired using Zen Blue software (Zeiss).

#### Experiment

Prior to imaging, cells were allowed to acclimate for at least 10min. After selection of scenes (see below), apoptotic cells were added to phagocyte-containing chamber slides and scenes were rapidly focused prior to the beginning of data collection. For each condition/experiment, at least 8 regions (‘scenes’) were selected and imaged. Scenes were selected if they featured at least 10 cells per region, and when experimental manipulations were performed (e.g., 6-AN treatment), regions were selected that featured similar cell numbers between vehicle and treatment conditions. Scenes were imaged every 4min for a minimum of 8h per experiment, which generally equates to ∼0.5μs per pixel or 5s per scene. Time-lapse experiments were performed in Multiplex CO-8Y mode which allows for gentle confocal-resolution imaging at a high frame rate. Specifically, we set the pinhole to 2.42 Airy units (57μm) with the image scaling (per pixel) set at 0.414μm x 0.414μm with 2x averaging. Focusing was ensured using the Definite Focus 2 module with a two-part strategy that was first based on the initial focus of individual scenes and then subsequently updated after each round of images. Analysis, including generation of scale bars, was performed using Zen Blue or Fiji.

### Ex Vivo Efferocytosis

To perform ex vivo efferocytosis, tissues were isolated from 7-week-old female C57BL/6 mice. To isolate bone marrow-resident macrophages (BMMs), bone marrow was flushed from bones using a 25G needle with α-MEM media containing FBS in a hypoxia chamber set to 1% oxygen. Red blood cells were lysed, and remaining bone marrow cells were plated in a non-TCtreated 6-well plate in BMDM media and cultured overnight in either hypoxia (1%) or standard (21%) oxygen. Floating cells were removed and fresh BMDM media was added. BMMs were then co-cultured with CypHer5E-labeled apoptotic MDA-MD-231 cells at a 1:1 ratio for 1h. Macrophages were isolated, stained with CD11b and F4/80, and analyzed via flow cytometry. To isolate splenic macrophages, spleens were minced into small pieces in a hypoxia chamber set to 1% oxygen and digested for 20min at 37C. Cell suspensions were strained using a 70μm cell strainer. Red blood cells were lysed, and remaining splenic cells were plated and treated identical to bone marrow-resident macrophages as described above.

### *In Vivo* Efferocytosis

To perform *in vivo* efferocytosis, we performed two models: dexamethasone (Dex)-induced apoptotic thymocyte removal and peritoneal apoptotic T cell removal. For Dex-induced clearance, six-to eight-week-old mice were injected i.p. with 300μl PBS containing 250μg dexamethasone (50μM; Sigma) with or without cycloheximide (5mg/kg) or 6-AN (3mg/kg) dissolved in EtOH. 6h after injection, thymi were harvested from mice and the numbers of thymocytes with annexin V staining only (apoptotic) versus annexin V/7-AAD double positive cells (secondarily necrotic) were assessed via flow cytometry. For the peritoneal clearance model, Cas9+ macrophages bearing either a scramble control guide or guides targeting G6pdx where initially conditioned in physiological hypoxia (1% O_2_) for 7d. Conditioned macrophages were subsequently i.p. injected 1h prior to injection of apoptotic cells. Then, mice were injected with 1e6 CypHer5E-labeled apoptotic Jurkat T cells and allowed to rest. Mice were subsequently euthanized 1h post-injection. Peritoneal lavage was collected in 8ml cold PBS. Collected lavage was centrifuged and cells were blocked with CD16/32 (clone 24G2) prior to staining with CD11b (eBioscience, 48-0112-82, clone M1/70, 1:750) and F4/80 (eBioscience, 12-4801-82, clone BM8, 1:750) for 30min at 4°C. Efferocytosis was analyzed by quantifying CypHer5E+ events within the CD11b+ F4/80high macrophages via flow cytometry.

### RNA Sequencing

For bulk RNA sequencing, 4.5×10^5^ macrophages were conditioned for 7d in standard O_2_ (∼21%) and either immediately collected or pulsed with 3h of 1% O_2_ (acute) and then collected. In parallel, 4.5×10^5^ macrophages were conditioned for 7d in 1% O_2_ (chronic) then collected. Collected samples were subsequently collected, washed with cold PBS, and lysed on ice with beta-mercaptoethanol-containing lysis buffer (Buffer RA, Machery-Nagel) for downstream RNA isolation. Total RNA was isolated using the NucleoSpin RNA isolation kit with on-column rDNase digestion (Machery-Nagel) and mRNA libraries were generated by polyA capture and reverse transcription of cDNA. Libraries were then sequenced at 150bp (paired-end) reads with ∼20 million reads per sample using an Illumina NovaSeq 6000 sequencer.

### Proteomics

For ^13^C SILAC labeling, target cells were grown in lysine- and arginine-deficient DMEM with 10% dialyzed FBS, 1% PSQ, and 2mM L-glutamine, supplemented with ‘heavy’ ^13^C_6_-lysine and ‘light’ ^12^C_6_-arginine (100mg/L; Cambridge Isotope Laboratories). After 8–10 passages, incorporation of ^13^C_6_-lysine-labeled amino acids into proteins was verified via LC-MS/MS to be >99%. Heavy isotope-labeled target cells were expanded in ^13^C_6_-lysine media and induced to undergo apoptosis. Macrophages were first conditioned in standard oxygen or hypoxia (see above), and then cultured with or without apoptotic cells for 1h. Apoptotic cells were subsequently removed using two cold assay media washes and two cold PBS washes. Macrophages were then removed using a cell scraper, pelleted to remove any residual liquid, and immediately snap frozen on dry ice.

#### Sample Preparation

Cell pellets were lysed with 200-300μL buffer containing 8M urea and 200mM EPPS (pH at 8.5) with protease inhibitor (Roche) and phosphatase inhibitor cocktails 2 and 3 (Sigma). Lysates were aspirated 2x on ice, followed by water sonication for 2min @4°C. Benzonase (Millipore) was added to a concentration of 50u/mL and incubated on ice for 15min. Samples were centrifuged at 14,000g for 10 min (@4°C) and the supernatant was subsequently extracted. The Pierce bicinchoninic acid (BCA) protein concentration assay was used to determine protein concentration. Protein disulfide bonds were reduced with 5mM tris (2-carboxyethyl) phosphine (@RT, 30min), then alkylated with 10mM iodoacetamide (@RT, 30min, in the dark). The reaction was quenched with 10mM dithiothreitol (@RT, 15min). Equivalent volumes of lysate aliquots were taken for each sample (100–200μg in each sample) and diluted to approximately 100μL with lysis buffer. Samples were subjected to chloroform/methanol precipitation as previously described (Navarrete-Perea et al., 2018). Pellets were reconstituted in 100μL of 200mM EPPS buffer and digested with Lys-C (1:100 enzyme-to-protein ratio) and incubated @37°C for 4h. Trypsin was then added (1:100 enzyme-to-protein ratio) and digested @37°C overnight. Anhydrous acetonitrile was then added to make a final volume of 30% ACN.

#### TMT Labeling

Samples were TMT-labeled as described (Navarrete-Perea et al., 2018). Briefly, samples were TMT-tagged by adding 10μL (28μg/μL) of TMTPro reagent for each sample and incubated for 1h @RT. A ratio check was performed by taking a 2μL aliquot from each sample and desalted by the StageTip method (Rappsilber et al., 2007) to confirm labeling efficiency. TMT-tags were then quenched with hydroxylamine to a final concentration of 0.3% for 15min @RT. Samples were pooled in their entirety, then dried via vacuum-centrifugation. Dried samples were reconstituted in 1mL of 3% ACN/1% TFA, desalted using a 100mg tC18 SepPak (Waters), and lyophilized overnight. Lyophilized peptides were dried using vacuum-centrifugation and reconstituted in 1mL of 2% ACN/25mM ABC. Peptides were fractionated into 48 fractions. Next, an Ultimate 3000 HPLC (Dionex) coupled to an Ultimate 3000 Fraction Collector using a Waters XBridge BEH130 C18 column (3.5μm x 4.6mm x 250mm) was operated at 1mL/min. The Buffers A, B, and C used below consisted of 100% water, 100% ACN, and 25mM ABC, respectively. The fractionation gradient operated as follows: 1% B to 5% B in 1min, 5% B to 35% B in 61min, 35% B to 60% B in 5min, 60% B to 70% B in 3min, 70% B to 1% B in 10min, with 10% C the entire gradient to maintain pH. The 48 fractions were then concatenated to 12 fractions (i.e., fractions 1, 13, 25, 37 were pooled, followed by fractions 2, 14, 26, 38, etc.) so that every 12th fraction was used to pool. Pooled fractions were vacuum-centrifuged then reconstituted in 1% ACN/0.1% FA for LCMS/MS.

#### LC-MS/MS

Total fractions were analyzed by LC-MS/MS using a NanoAcquity (Waters) with a 50cm EASY-Spray Column (PepMap RSLC, C18, 2µm, 100Å, 75µm I.D.) heated to 60°C coupled to a Orbitrap Eclipse Tribrid Mass Spectrometer (Thermo Fisher Scientific). Peptides were separated by direct inject at a flow rate of 300nL/min using a gradient of 5 to 30% acetonitrile (0.1% FA) in water (0.1% FA) over 3h then to 50% ACN in 30min and analyzed by SPS-MS3. MS1 scans were acquired over a range of m/z 375-1500, 120K resolution, AGC target (4.0e5), and maximum IT of 50ms. MS2 scans were acquired on MS1 scans of charge 2-7 using an isolation of 0.7m/z, collision induced dissociation with activation of 32%, turbo scan and max IT of 50ms. MS3 scans were acquired using specific precursor selection (SPS) of 10 isolation notches, m/z range 100-1000, 50K resolution, AGC target (1.0e5), HCD activation of 45%, and max IT of 150ms. The dynamic exclusion was set at 60s.

#### TMT Data Analysis

Raw data files were processed using Proteome Discoverer (PD) version 2.4.1.15 (Thermo Scientific). For each of the TMT experiments, raw files from all fractions were merged and searched with the SEQUEST HT search engine with a UniProt protein database downloaded on 2019/01/09 (176,945 entries). The precursor and fragment mass tolerances were 10ppm and 0.6Da respectively. A maximum of two trypsin missed cleavages were permitted. Searches used a reversed sequence decoy strategy to control peptide false discovery rate (FDR) and 1% FDR was set as threshold for identification.

### Analysis of Nutrient Consumption and Secretion via YSI

Macrophages (2.5×10^6^) were cultured in either standard oxygen or hypoxia (1% O_2_) and conditioned for a total of 7 days. Media was changed at day 1, day 3, and day 5 prior to a 24h period of conditioning. At the end of the 24h period (denoted as day 2-3, day 4-5, and day 6-7 periods), media was collected, spun down to remove cell debris, and immediately snap frozen for downstream analysis via a 2950D Biochemistry Analyzer (YSI Life Sciences) to determine glucose and lactate concentrations. Absolute rates of consumption (glucose) or secretion (lactate) were calculated first by subtracting the concentration observed in control (cell-free) media incubated in parallel, then normalized to cell number and volume of media. These experiments were performed independently at least two times.

### Untargeted Metabolomics

Macrophages (2.5×10^6^) were conditioned in standard oxygen or hypoxia for 7d. To protect from experimental artifact, all samples were collected in the same oxygen environment that they were conditioned in. Plates containing cells were placed on ice and washed with cold PBS three times, then moved to dry ice and ice cold 80% methanol was added. Cells were subsequently scraped and transferred to a cold centrifuge tube on dry ice. The cell-methanol slurries were then vortexed for 1min, allowed to rest on dry ice for 5min, then vortexed an additional 1min to create cell lysates. Lysates were centrifuged for 20min at 14,000g in a refrigerated centrifuge set to 4°C to remove debris. Supernatants were transferred to clean tubes and lyophilized in the absence of heat and dissolved in water. Targeted LC/MS analyses were performed on a Q Exactive Orbitrap mass spectrometer (Thermo Scientific) coupled to a Vanquish UPLC system (Thermo Scientific). The Q Exactive operated in polarity-switching mode. A Sequant ZIC-HILIC column (2.1mm i.d. × 150mm, Merck) was used for separation of metabolites. Flow rate was set at 150μL/min. Buffers consisted of 100% acetonitrile for mobile B, and 0.1% NH4OH/20mM CH3COONH4 in water for mobile A. Gradient ran from 85% to 30% B in 20min followed by a wash with 30% B and re-equilibration at 85% B. MS data were processed using Compound Discoverer (Thermo Scientific). An in-house metabolite library as well as Chemspider were searched for metabolite identification. Three levels of metabolite identification were reported: 1) identified compounds: definitive identification based on the mass (within 5ppm) and retention time of authentic chemical standards; 2) putatively annotated compounds by searching ChemSpider (mass tolerance 5ppm); 3) compounds with predicted chemical composition based on mass. Relative metabolite quantitation was performed based on peak area for each metabolite. Hierarchical clustering analysis and principal component analysis (PCA) were performed by Compound Discoverer (Thermo Scientific).

### 13C-Metabolic Flux Analysis

Experiments were performed similar to those outlined in Untargeted Metabolomics with the following modification. On day 5, glucose-deficient RPMI containing 1 g/L [1,2-^13^C]glucose or [U-^13^C]-glucose (both from Cambridge Isotope Laboratories), 10% L929-conditioned media, 1% PSQ, and 10% FBS was conditioned in either standard oxygen or hypoxia for 16-24h. On day 6, conditioned macrophages were washed with PBS twice and then cultured with heavy isotope-containing media for 16h. Metabolites were extracted using the same protocol detailed in Untargeted Metabolomics. LC/MS analyses were performed on a Q Exactive Orbitrap mass spectrometer (Thermo Scientific) coupled to a Vanquish UPLC system (Thermo Scientific). The Q Exactive operated in polarity-switching mode. A Sequant ZIC-HILIC column (2.1mm i.d. × 150mm, Merck) was used for separation of metabolites. Flow rate was set at 150μL/min. Buffers consisted of 100% acetonitrile for mobile B, and 0.1% NH4OH/20mM CH3COONH4 in water for mobile A. Gradient ran from 85% to 30% B in 20min followed by a wash with 30% B and re-equilibration at 85% B. Data analysis was done using El-MAVEN (v0.12.0). Metabolites and their ^13^C isotopologues were identified on the basis of exact mass within 5ppm and standard retention times. Relative metabolite quantitation was performed based on peak area for each metabolite.

### Analysis of Mitochondrial Properties

For analysis of mitochondrial superoxide levels, conditioned macrophages were labeled with the live cell-permeant fluorogenic probe MitoSOX Red (ThermoFisher). MitoSOX Red is targeted to the mitochondria and is oxidized by superoxide but not reactive oxygen or nitrogen species, inducing an increase in fluorescence which we quantified using flow cytometry. To estimate mitochondrial shape and content, conditioned macrophages were labeled with both MitoTracker Green (MTG) and MitoTracker Deep Red (MTDR, both ThermoFisher). MTG localizes to mitochondria independent of mitochondrial membrane potential whereas MTDR labeling of mitochondria is dependent on mitochondrial membrane potential. Combined, the two fluorogenic probes were used to estimate potential and quantity (via flow cytometry).

### NADP+/NADPH, ATP, and GSH/GSSG Measurement

4×10^6^ or 2.5×10^4^ (for GSH/GSSG) macrophages were seeded for conditioning and subsequent NADP+/NADPH, ATP, or GSH/GSSG measurement. In some experiments, macrophages were treated with 6-AN (100nM) or vehicle (DMSO) beginning one day after replating. On day 7, cells were collected, processed, and the ratio of NADP+/NADPH or GSH/GSSG was measure using the NADP/NADPH Quantitation Colorimetric Kit (BioVision) or the GSH/GSSG-Glo Assay (Promega), respectively, or ATP was quantitated using the Luminescent ATP Detection Assay Kit (Abcam), following the manufacturer’s protocol.

### Measurement of Redox State and Lysosomal Acidity

Lipid peroxidation and cellular ROS was determined by labeling of phagocytes using the reagent BODIPY 581/591 C11 and CellROX Deep Red (both ThermoFisher), respectively, followed by flow cytometric analysis. Lysosomal acidity was determined by labeling of phagocytes using the reagent Lysosensor Green DND-189. Labeling was performed according to manufacturer instructions. For analysis of redox state or lysosomal acidity in efferocytotic cells, phagocytes were first gated on CypHer5E+.

### CRISPR/Cas9 Deletion or siRNA knockdown of SLCs

Stable, individual clones of Cas9-expressing Hoxb8 macrophages were generated using bone marrow from Cas9 mice (JAX Strain #026179) and a lentiviral transduction protocol adapted from Fang Zhang and colleagues. *G6pdx* was disrupted using the Zhang lab lentiGuide-Puro sgRNA plasmid with a verified guide for *G6pdx*. lentiGuide-Puro was a gift from Feng Zhang (Addgene plasmid # 52963). Guide RNAs targeting *G6pdx* and scramble control were generated using the following oligo pairs:

G6pdx:

5’---CACCGACCTGAAGATACCTTCATTG---3’

5’---AAACCAATGAAGGTATCTTCAGGTC---3’

Scramble Control:

5’---CACCGCACTCACATCGCTACATCA---3’

3’---AAACTGATGTAGCGATGTGAGTGC---5’.

### Statistics and Reproducibility

Statistical analyses were executed using GraphPad Prism 7, SPSS v.22, and R v.4.0.5. We determined statistical significance, depending on the structure of the data, via unpaired twotailed Student’s t-test, nonparametric Mann–Whitney U-test, one-way or two-way ANOVA, or Fisher’s exact test. R v.4.0.5 was used for graphical and statistical analyses and the R package DESeq2 was used for differential gene expression analysis of transcriptomic and proteomic analyses. All genes were curated according to a previously described approach (Morioka et al., 2018). Complementary analyses of biological pathways were performed by comparing significantly differentially expressed genes or proteins and cross-referencing them with the Molecular Signatures Database (MSigDB). Gephi (v.0.9.1 https://gephi.org/) was used to perform standard network clustering analyses. The “Link Communities” algorithm for biological network analysis was used to calculate edges and nodes (Ahn et al., 2010). The Yifan Hu layout algorithm was used to determine network structure. All biologically independent samples were included and used for statistical and graphical analyses. No data was excluded from this manuscript. Sample sizes were not predetermined using statistical methods. The code used in this manuscript is available upon reasonable request. RNA sequencing data is deposited under NCBI GEO accession GSE192969.

## Acknowledgements

We thank members of the Perry laboratory and co-authors for edits and discussions related to this manuscript. We thank Justin Cross and the Donald B. and Catherine C. Marron Cancer Metabolism Center at MSKCC for help with YSI experiments. This work was supported by grants to J.S.A.P. from the NIH (NCI 5R00CA237728; NIGMS 1DP2GM146337), a Parker Institute for Cancer Immunotherapy Career Development Award, a V Foundation Scholars Award, a MSKCC Functional Genomics Institute pilot award, a grant to K.R.K. from the NIH (NCI 1R01CA248364), and MSKCC Cancer Center Support Grant P30CA008748. Some figure panels were created using the commercial version of BioRender.

## Competing Financial Interests

K.R.K. Serves on the scientific advisory board of NVision Imaging Technologies. He holds patents related to imaging and modulation of cellular metabolism.

