## Supplemental Figures for "Novel adaptation supports enhanced macrophage efferocytosis in limited-oxygen environments"

### Supplemental Figures 1 – 8

**Figure S1: Efferocytosis is enhanced under prolonged ('chronic') physiological hypoxia**

(A) UV-induced apoptosis in different targets. Apoptosis was induced in MDA-MB-231s, BrM2s, and Jurkat T cells via UV-C irradiation and monitored for cell death at the time points indicated. (Left) Representative FACS plots of apoptosis as assessed using Annexin V (AV) and 7AAD. (Right) Quantification of $AV^{+}7AAD^{+}$, $AV^{+}$, $7AAD^{+}$, $DN^{+}$ (double negative) at different time points for each target cell. Data represent three experimental replicates. Data shown as mean ± SEM.

(B) Efferocytosis by standard oxygen- or hypoxia-conditioned macrophages of primary thymocytes treated with dexamethasone. Standard or chronic hypoxia-conditioned ER-Hoxb8 macrophages as in (**Figure 1A**) were co-cultured with CypHer5E-labeled apoptotic primary thymocytes at a 1:1 ratio. Instead of UV-C, cell death was induced using dexamethasone. Efferocytosis was assessed via FACS. Data are from four independent experiments. Data are shown as mean ± SEM. ****p < .0001.

(C, D) Efferocytosis by standard oxygen- or hypoxia-conditioned bone marrow-derived macrophages of primary thymocytes treated with UV-C or dexamethasone. Standard or chronic hypoxia-conditioned bone marrow-derived macrophages were co-cultured with CypHer5E-labeled apoptotic primary thymocytes at a 1:1 ratio. Cell death was induced using UV-C (C) or dexamethasone (D). Efferocytosis was assessed via FACS. Data are from four independent experiments. Data are shown as mean ± SEM. ****p < .0001.

(E) FACS assessment of efferocytotic capacity. Conditioned macrophages were co-cultured with TAMRA-labeled apoptotic MDA-MB-231 cells at a 1:1 ratio for 1h and analyzed for efferocytosis. (Left) TAMRA+ percentage and (Right) TAMRA geometric mean of efferocytotic standard oxygen- and chronic hypoxia-conditioned macrophages. Data represent three independent experiments. Data shown as mean ± SEM. ****p < .0001; ns = not significant.

**Figure S1: Efferocytosis is enhanced under prolonged ('chronic') physiological hypoxia**

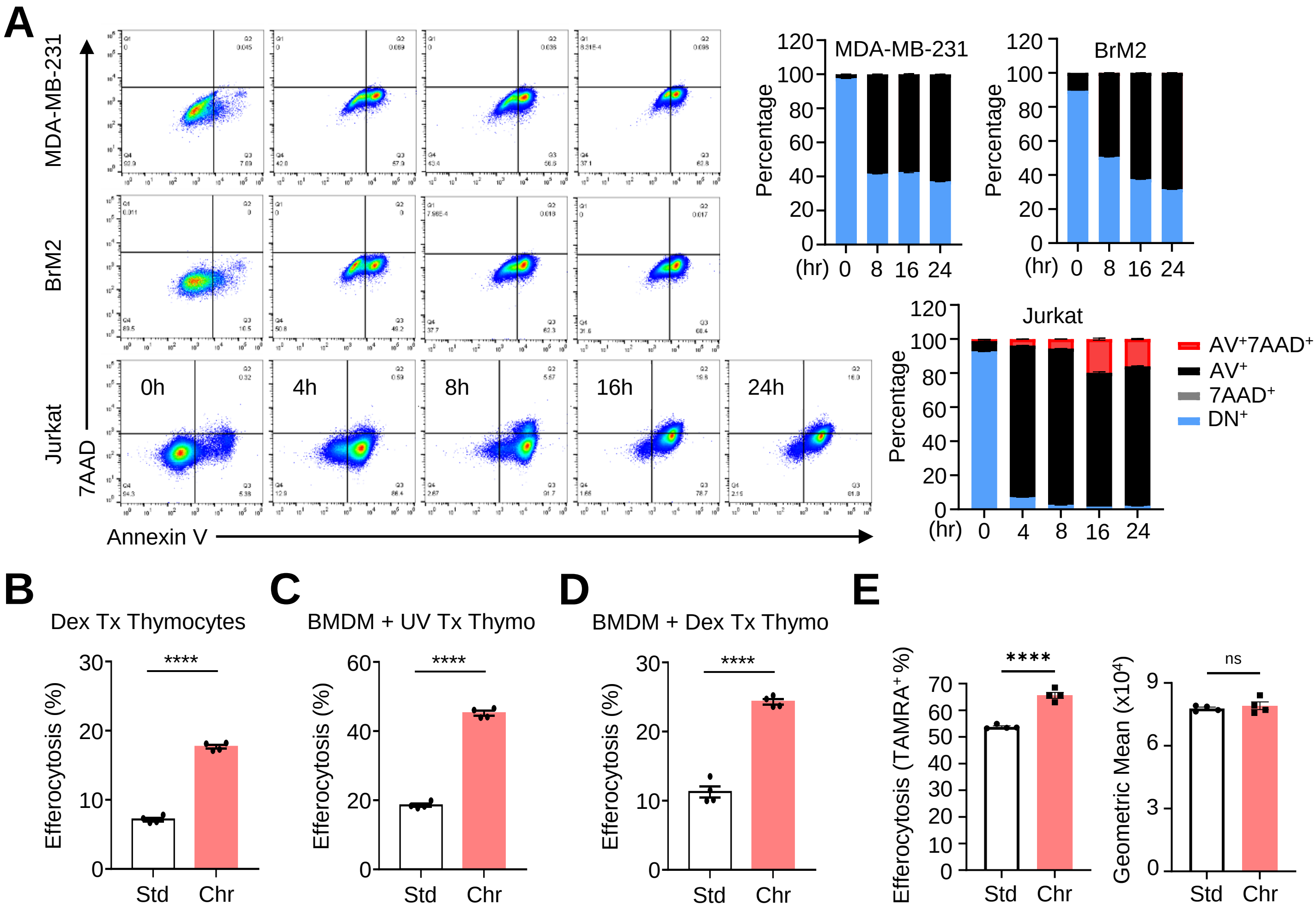

**Figure S2: Characterization of macrophages under chronic physiological hypoxia**

(A, B) Functional programs and wound healing transcripts differentially regulated in chronic physiological hypoxia-conditioned macrophages. (A) Additional analyses of RNA sequencing data obtained, analyzed, and presented as in **Figure 2A**. (B) Expression of known wound healing and pro-inflammatory macrophage-associated genes in standard oxygen- and chronic physiological hypoxia-conditioned macrophages analyzed via RNA sequencing as in (**Figure 2**). A bolded transcript name denotes a gene that is significantly differentially regulated by false discovery rate-corrected statistical analysis. *Il4ra* - *p* value = .054. *Tnf* - *p* value = .094. *Itgam* - *p* value = .079.

**Figure S2. Characterization of macrophages under chronic physiological hypoxia**

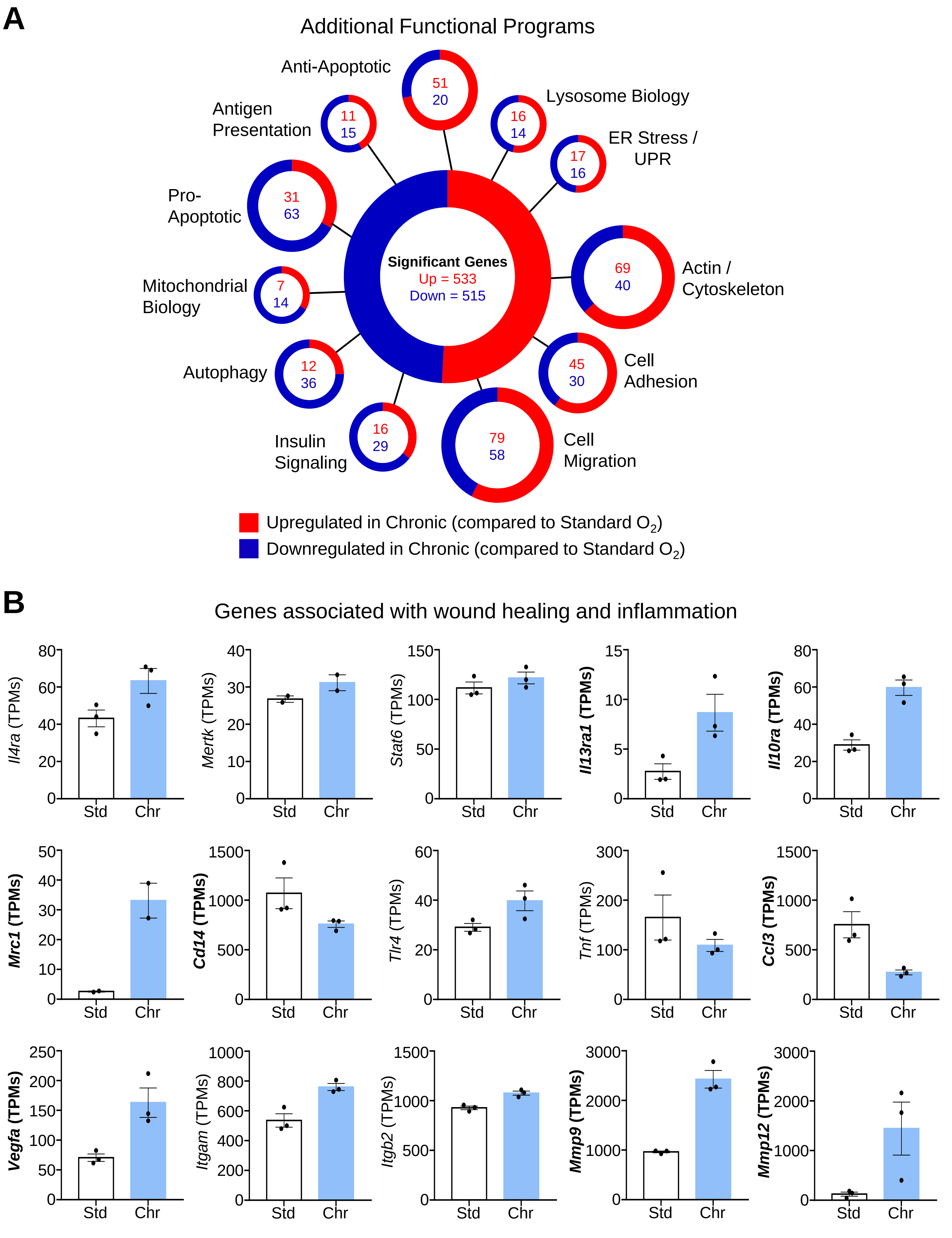

**Figure S3: Chronic hypoxia induces a metabolic state poised for efficient efferocytosis**

(A) Analysis of chronic physiological hypoxia-conditioned macrophage transcripts also found via proteomics. Shown is the cross-reference analysis of RNA sequencing data from **Figure 2A** and global proteomic data from **Figure 2D**. (Top) Upregulated transcripts not detected in proteomics (orange), detected and found significantly upregulated in proteomics (red), or detected but found not significantly upregulated in proteomics (gray). (Bottom) Downregulated transcripts not detected in proteomics (purple), detected and found significantly downregulated in proteomics (blue), or detected but found not significantly downregulated in proteomics (gray).

(B) Global proteomic analysis of resting and efferocytotic macrophages in standard oxygen versus chronic physiological hypoxia conditions. Shown are the number of differentially up- or downregulated proteins in comparisons between naïve and efferocytotic standard oxygen-conditioned macrophages (Top Left), between naïve and efferocytotic chronic physiological hypoxia-conditioned macrophages (Top Right), and between efferocytotic standard oxygen and physiological hypoxia-conditioned macrophages (Bottom).

(C) Functional program analysis of resting and efferocytotic macrophages in standard oxygen versus chronic physiological hypoxia conditions. Similar to analyses in **Figure 2D**, shown are functional program analyses of differentially expressed proteins from comparisons shown in Figure S3B.

**Figure S3: Chronic hypoxia induces a metabolic state poised for efficient efferocytosis**

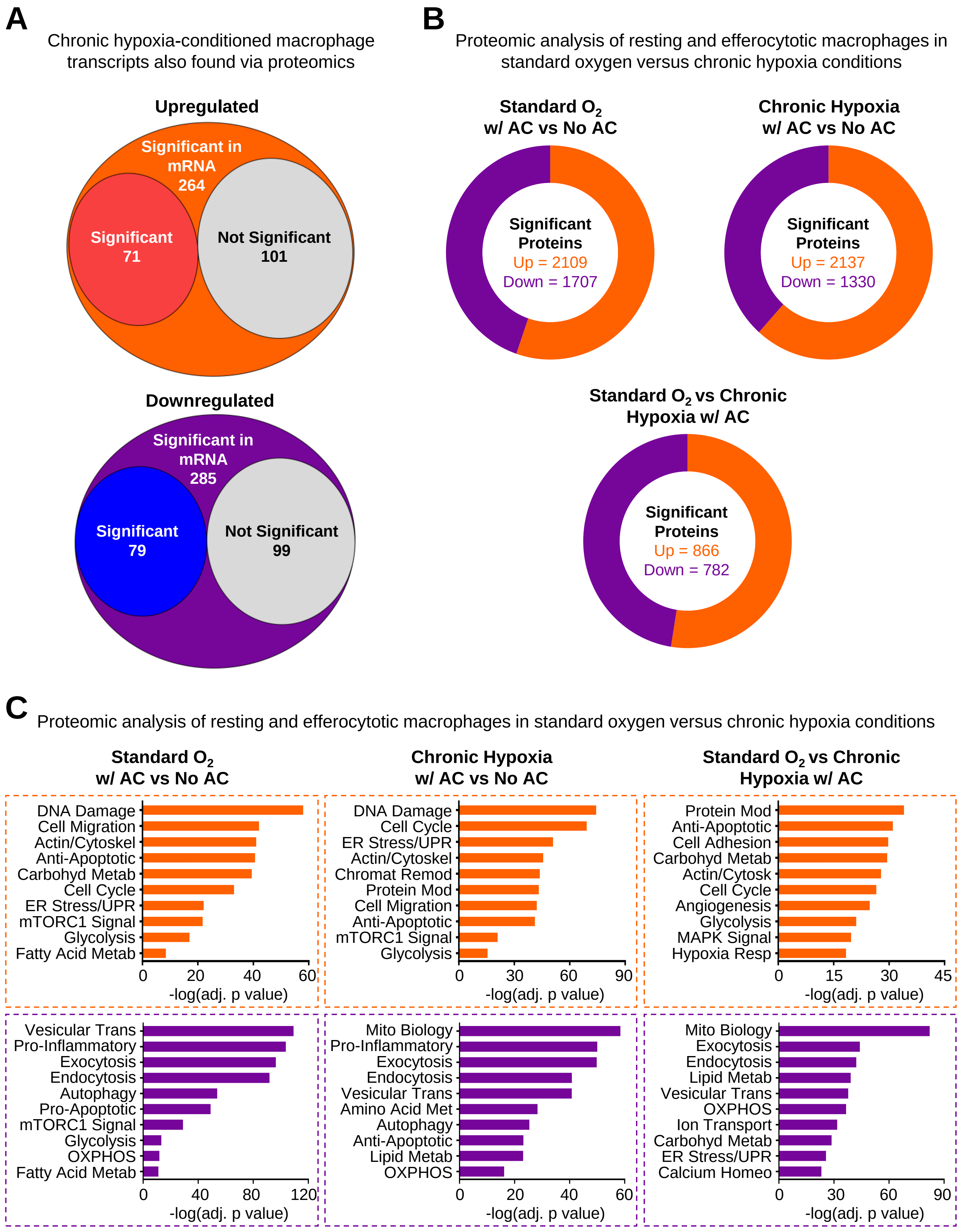

**Figure S4: Chronic hypoxia induces a metabolic state poised for efficient efferocytosis**

Representative genes/proteins of 'primed' and 'poised' states. Shown are additional representative transcripts/proteins from each state observed in **Figure 3A and 3B**. PGAM1/*Pgam1*, Phosphoglycerate mutase 1. GAPDH/*Gapdh*, Glyceraldehyde-3-phosphate dehydrogenase. TPI/*Tpi*, Triosephosphate isomerase. ALDOA/*Aldoa*, Fructose-bisphosphate aldolase A. PGM2/*Pgm2*, Phosphopentomutase. AK4/*Ak4*, Adenylate kinase 4. MAN1A/*Man1a*, Mannosyl-oligosaccharide 1,2-alpha-mannosidase IA. EGLN1/*Egln1*, Egl nine homolog 1. PTPRO/*Ptpro*, Receptor-type tyrosine-protein phosphatase O. PLOD1/*Plod1*, Procollagen-lysine,2-oxoglutarate 5-dioxygenase 1.

**Figure S4: Chronic hypoxia induces a metabolic state poised for efficient efferocytosis**

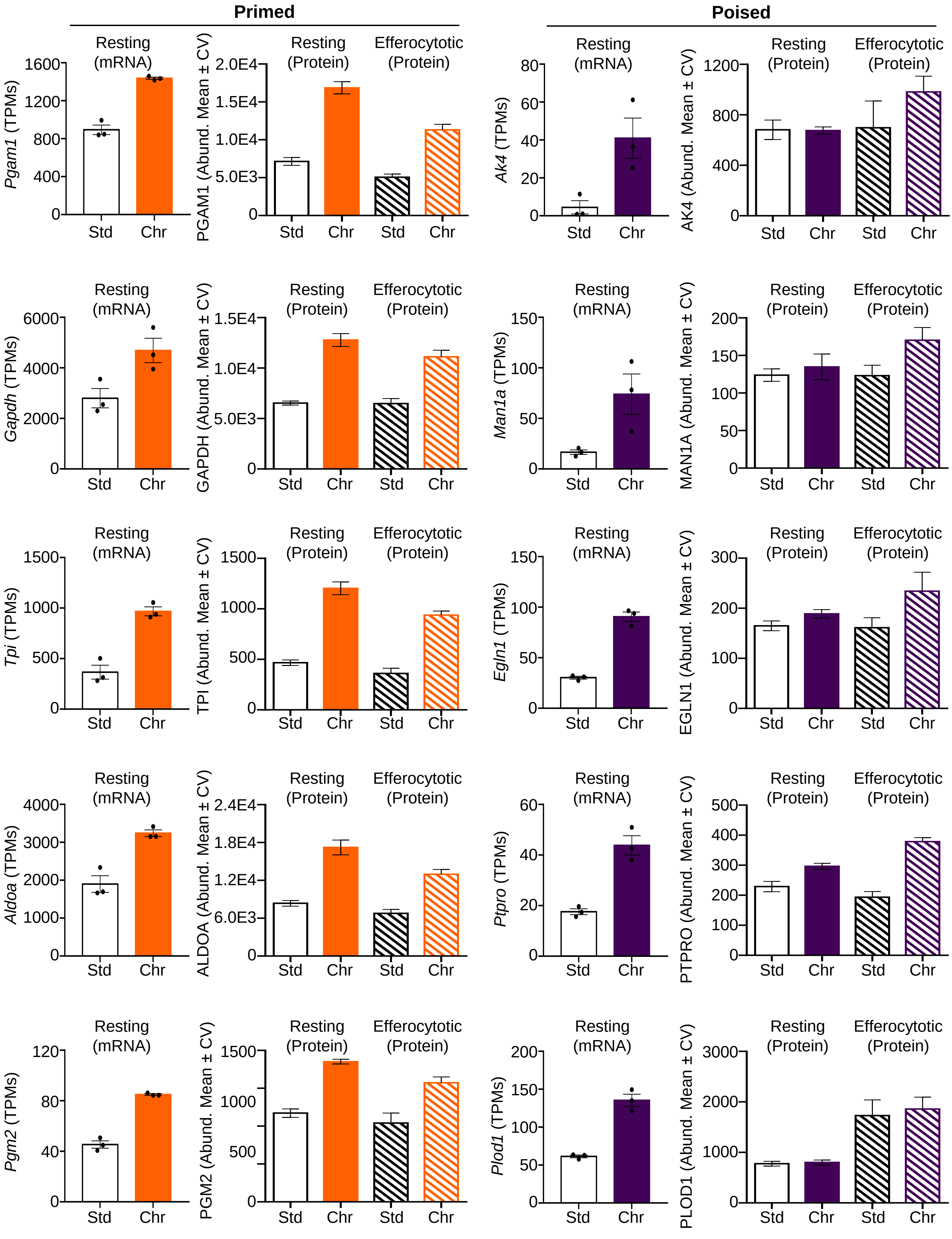

**Figure S5: Mapping metabolic pathway use in chronic hypoxia-conditioned macrophages**

(A) Network analysis of differentially expressed metabolic transcripts identified in chronic physiological hypoxia-conditioned macrophages. Analyses performed as in **Figure 4A**. Metabolic genes that are differentially regulated in chronic physiological hypoxia-conditioned macrophages (**Figure 2**) are represented using network analysis to determine family clusters (shaded areas) and connectedness between individual genes.

(B) Principal components analysis of untargeted metabolomics analysis of standard oxygen and physiological hypoxia-conditioned macrophages. Shown is a principal components analysis (PCA) of experiments performed in **Figure 4D**. Principal component (PC)1, which accounts for 74.3% of the total variance, is entirely accounted for by chronic physiological hypoxia-conditioned macrophages, whereas PC2 (12.4% of total variance) represents the acute physiological hypoxia vs standard oxygen comparison.

(C) Representative individual metabolites from untargeted metabolomics analysis. As in **Figure 4E**, shown are representative metabolites from significantly differentially regulated pathways from **Figure 4D**, specifically upper glycolysis (blue), pentose phosphate pathway (purple), lower glycolysis/TCA cycle (orange), and alanine/aspartate/glutamine metabolism (gray). Data are shown as mean ± SEM. All metabolites except for pyruvate are significant, ranging from $p < .0001$ to $p < .05$. F6P, fructose 6-phosphate. FBP, fructose 1,6-bisphosphate. R5P, ribose 5-phosphate. SAM, S-adenosyl methionine. SAH, S-adenosyl homocysteine. Glyc3P, glycerol 3-phosphate.

**Figure S5: Metabolic pathway use in chronic physiological hypoxia-conditioned macrophages**

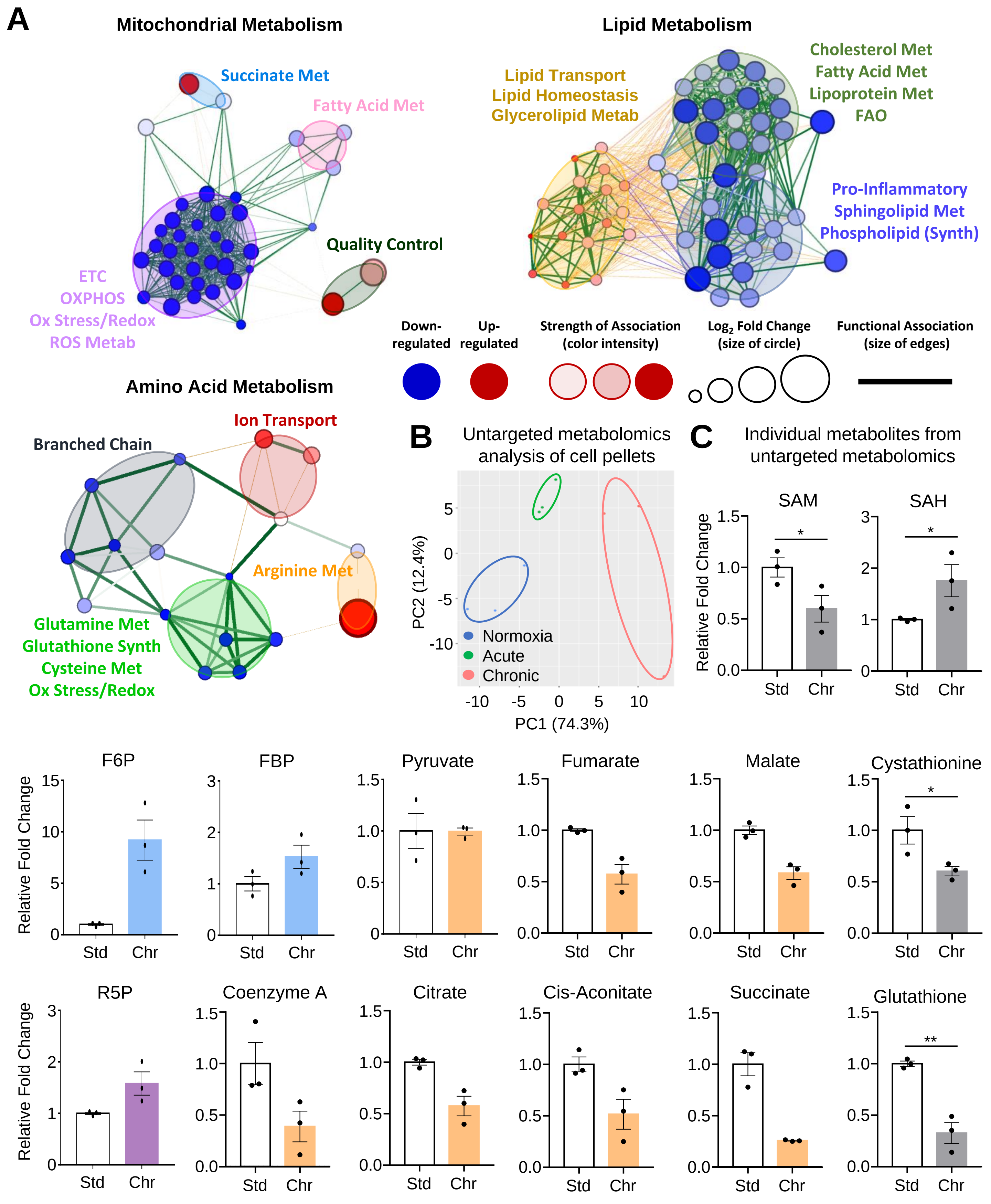

**Figure S6: Macrophages co-opt noncanonical pentose phosphate loop for redox homeostasis**

(A) The pentose phosphate pathway is active in chronic physiological hypoxia-conditioned macrophages. (Left) Data from experiments performed in **Figure 5A**. Shown is the relative abundance of the indicated isotopologues. Intermediates from PPP are denoted in red, gluconeogenesis denoted in green, conventional glycolysis denoted in blue, and TCA cycle denoted in orange. For fractional enrichment, see **Figure 5A**. Three independent experiments were performed for each condition. Data shown as mean ± SEM. *p < .05; **p < .01; ***p < .001; ****p < .0001. G6P, glucose 6-phosphate. F6P, fructose 6-phosphate. FBP, fructose 1,6-bisphosphate. 3PG, 3-phosphoglyceric acid. PEP, phosphoenolpyruvate. S7P, sedoheptulose 7-phosphate.

(B) Identification of flux through novel pentose phosphate pathway intermediates in chronic hypoxia-conditioned macrophages. Data from experiments performed in **Figure 5A**. Shown is the fractional enrichment of the indicated isotopologues from **Figure 5C**. Gluconeogenesis intermediates are denoted in green. Three independent experiments were analyzed for each condition. Data shown as mean ± SEM. *p < .05; **p < .01; ***p < .001; ****p < .0001; ns = not significant. G6P, glucose 6-phosphate. S7P, sedoheptulose 7-phosphate. F6P, fructose 6-phosphate. FBP, fructose 1,6-bisphosphate. 3PG, 3-phosphoglyceric acid.

(C) Gluconeogenesis is prevalent in chronic physiological hypoxia-conditioned macrophages. Data from experiments performed in **Figure 5D**. Shown is the fractional enrichment of the indicated isotopologues from **Figure 5D**. PPP intermediates are denoted in red, gluconeogenesis in green, and conventional glycolysis in blue. Three independent experiments were performed for each condition. Data are shown as mean ± SEM. *p < .05; ***p < .001.

(D) Less glucose is converted to alanine in chronic physiological hypoxia-conditioned macrophages. Abundance (Left) and labeled fraction (Right) of M+3 alanine from experiments performed in **Figure 5D**. Three independent experiments were performed for each condition. Data are shown as mean ± SEM. ***p < .001; ns = not significant.

(E) Less glucose enters the TCA cycle in chronic physiological hypoxia-conditioned macrophages. Data are from experiments performed in **Figure 5D**. Glucose entry into the TCA cycle can occur via citrate (M+2) or oxaloacetate (M+3). Shown is the relative abundance of the indicated isotopologues. For fractional enrichment, see Figure S5E. Three independent experiments were performed for each condition. Data shown as mean ± SEM. *p < .05; **p < .01.

(F) Less glucose contributes to nucleotide synthesis in chronic physiological hypoxia-conditioned macrophages. The $^{12}C$ and $^{13}C$ abundance of IMP (Top) and GMP (Bottom) from experiments performed in **Figure 5D**. Three independent experiments were performed for each condition. Data are shown as mean ± SEM. *p < .05; **p < .01; ***p < .001.

### Figure S6: Macrophages co-opt noncanonical pentose phosphate loop for redox homeostasis

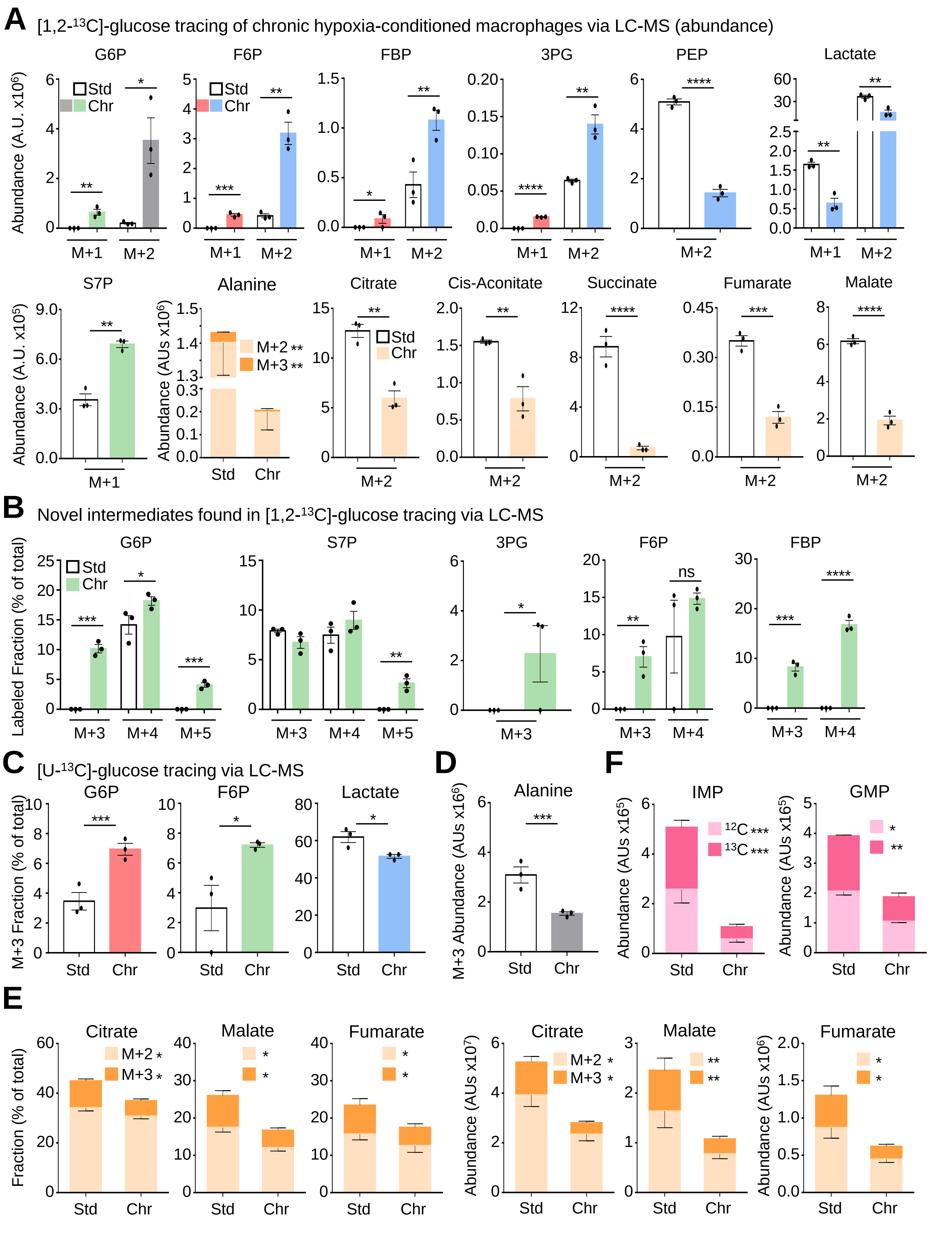

**Figure S7: Macrophages co-opt noncanonical pentose phosphate loop for redox homeostasis**

(A) NADP$^+$ level and NADP+/NADPH ratio in standard oxygen and chronic physiological hypoxia-conditioned macrophages. Data are from experiments performed as in **Figure 5F**. Three independent experiments were performed for each condition. Data are shown as mean ± SEM. ****p < .0001. ns = not significant.

(B, C) Mitochondrial membrane potential (MMP) and mitochondrial mass of conditioned macrophages. Macrophages were differentiated and conditioned as in **Figure 1A** and labeled with Mitotracker Deep Red (B for MMP) or Mitotracker Green (C for mitochondrial mass). Macrophages were subsequently analyzed by flow cytometry. For FACS analysis, plots represent two biological experiments which are summarized as the MFI fold change relative to standard oxygen-conditioned macrophages. Data shown as mean ± SEM.

(D) Mitochondrial ROS levels in conditioned macrophages. Macrophages were differentiated and conditioned as in **Figure 1A**, labeled with MitoSox Red, and analyzed via flow cytometry. Shown are representative FACS plots (Left) and summary plots of the MFI fold change relative to standard oxygen-conditioned macrophages (Right). Data are representative of three independent experiments per condition. Data are shown as mean ± SEM. ****p < .0001.

(E) Total glutathione, GSH, and GSSG in conditioned macrophages. Macrophages were differentiated and conditioned as in **Figure 1A**. Cells were subsequently analyzed via plate reader. Net relative luminescence unit (RLU) values were normalized to cell number. Data are from four independent experiments for each condition. Data are shown as mean ± SEM. **p < .01; ***p < .001. See also **Figure 5H**.

**Figure S7: Macrophages co-opt noncanonical pentose phosphate loop for redox homeostasis**

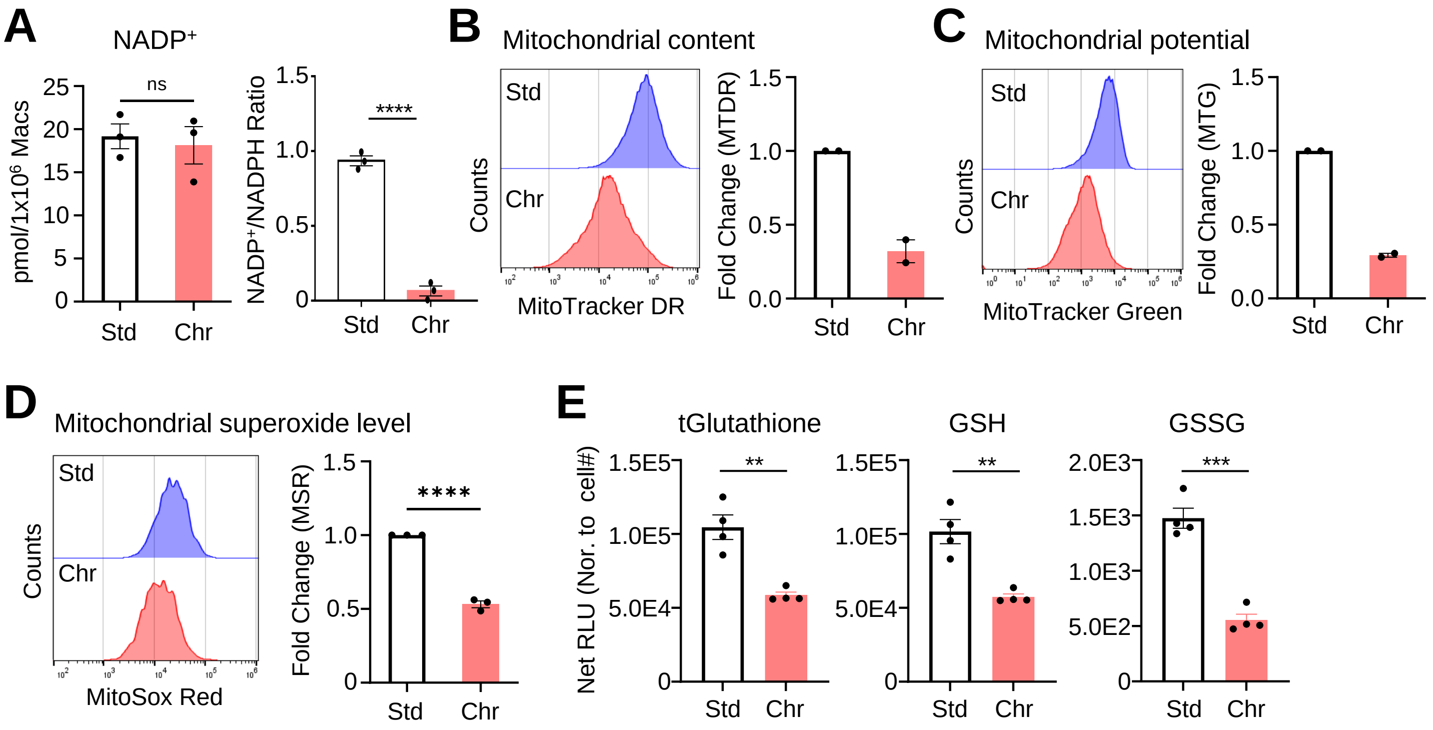

**Figure S8: Noncanonical pentose phosphate loop supports continual efferocytosis**

(A, B) Targeting PPP metabolism or gluconeogenesis reduces efferocytosis specifically in chronic hypoxia-conditioned macrophages. (A) Schematic illustrating the approaches used to target key PPP enzymes (G6PDi targeting G6PD and 6-AN targeting 6PGD) and gluconeogenesis (FBPi targeting FBP1 and exogenous lactate). (B) Macrophages were differentiated and conditioned as in **Figure 1A**. Indicated treatments (or vehicle controls) were added prior to the start of an assays. Macrophages were subsequently cultured with CypHer5E-labeled apoptotic MDA-MB-231s at a 1:1 ratio for 1h in the presence of indicated inhibitors (or vehicle controls). Efferocytosis was assessed via flow cytometry. Data are representative of three independent experiments. Data are shown as mean ± SEM. **p < .01; ***p < .001; ****p < .0001; ns = not significant.

(C) Western blot analysis of G6PDX. Cas9/GFP+ Hoxb8 macrophages bearing scramble (WT) or *G6pdx* (G6) small guides were analyzed for G6PDX and GAPDH protein levels.

(D) Lysosomal acidity increases in chronic hypoxia-conditioned efferocytotic macrophages with disrupted PPP metabolism. Shown are representative FACS plots from **Figure 7D**.

(E) Lipid peroxidation increases in chronic hypoxia-conditioned efferocytotic macrophages with disrupted PPP metabolism. Macrophages were differentiated and conditioned as in **Figure 6B**. Conditioned macrophages were cultured with CypHer5E$^{+}$-labeled apoptotic MDA-MB-231 cells at a 1:1 ratio for 1h. Uncleared apoptotic cells were removed, and macrophages were stained with C11-BODIPY 581/591. CypHer5E$^{+}$ macrophages were gated as actively efferocytotic for analysis of lipid peroxidation. Shown are representative FACS plots (Left) and MFI (Right) from three independent experiments per condition. Data are shown as mean ± SEM. *p < .05; ***p < .001; ****p < .0001.

(F) Cellular ROS increases in chronic hypoxia-conditioned efferocytotic macrophages with disrupted PPP. Shown are representative FACS plots from **Figure 7F**.

**Figure S8: Noncanonical pentose phosphate loop supports continual efferocytosis**

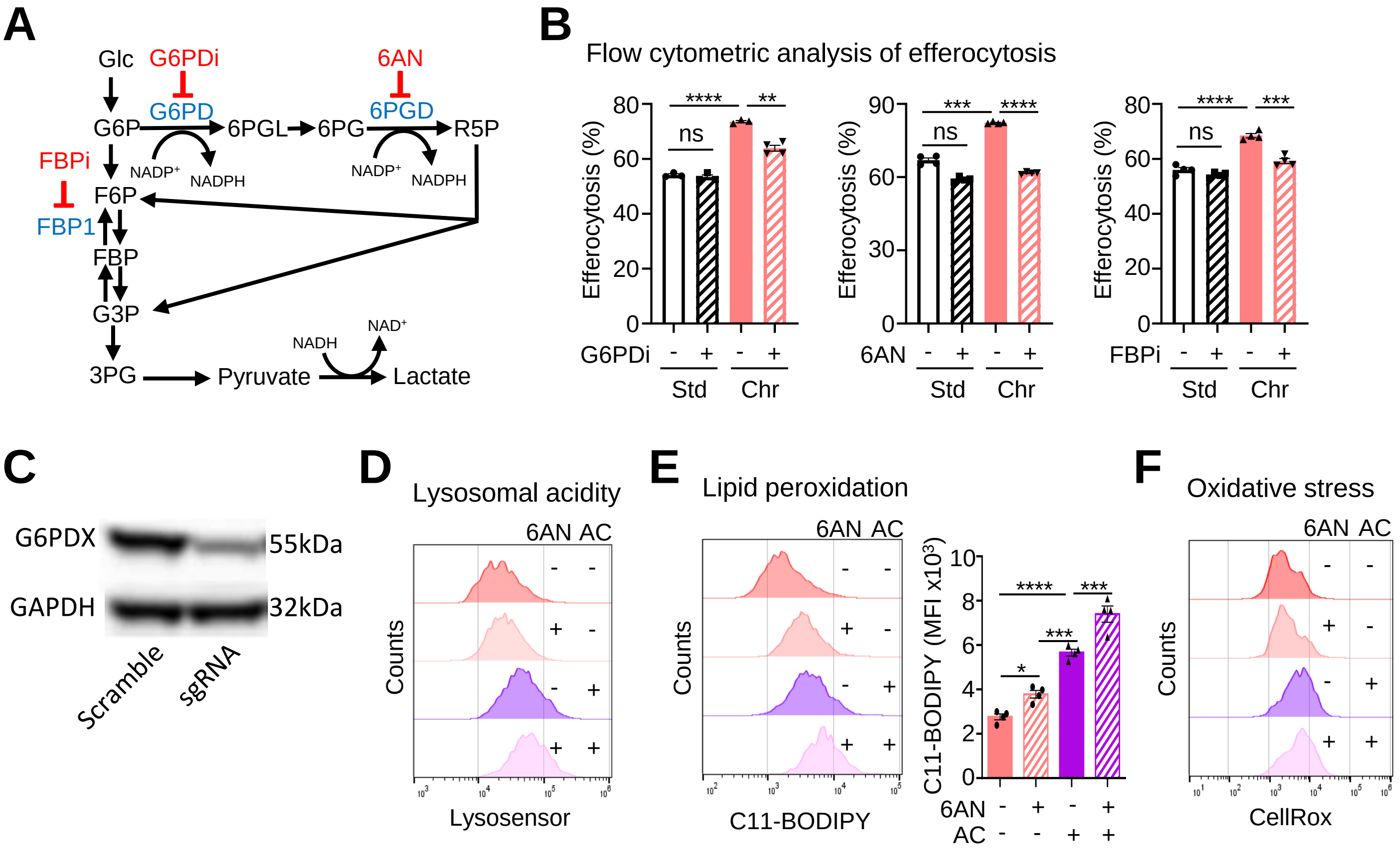
